# Scale-dependent effects of geography, host ecology, and host genetics, on species composition and co-occurrence in a stickleback parasite metacommunity

**DOI:** 10.1101/672410

**Authors:** Daniel I. Bolnick, Emlyn J. Resetarits, Kimberly Ballare, Yoel E. Stuart, William E. Stutz

## Abstract

A core goal of ecology is to understand the abiotic and biotic variables that regulate species distributions and community composition. A major obstacle is that the rules governing species distribution can change with spatial scale. Here, we illustrate this point using data from a spatially nested metacommunity of parasites infecting a metapopulation of threespine stickleback fish from 34 lakes on Vancouver Island, British Columbia. Parasite communities differ among host individuals within each host population and between host populations. The distribution of each parasite taxon depends, to varying degrees, on individual host traits (e.g., mass, diet) and on host population characteristics (e.g., lake size, mean diet). However, in most cases, a given parasite was regulated by different factors at the host-individual and host-population scales, contributing to scale-dependent patterns of parasite-species co-occurrence.

## Introduction

A classic dichotomy in ecology is whether communities are deterministic co-occurring sets of species (Clements 1916) or collections of many species following an independent set of stochastic rules (Gleason 1926). Metacommunity theory (Leibold et al. 2004, Leibold and Chase 2017) bridges the gap between these competing visions by considering relative roles of determinism and stochasticity at various spatial scales on a fragmented landscape. When species have similar filters governing dispersal to new patches and persistence within patches, they will tend to co-occur and form a more Clementsian community. If, instead, each species’ distribution is stochastic, or is subject to species-specific filters, then communities will be more Gleasonian. Thus, a key question in metacommuity theory is, what abiotic and biotic filters regulate species’ dispersal or within-patch dynamics? Then, do these filters affect multiple species in a similar manner? Here, we present a case study using a multi-species metacommunity of parasites.

Parasite communities are an ideal system to apply metacommunity theory (Lima et al. 2012, Mihaljevic 2012, Seabloom et al. 2015, Borer et al. 2016). Metacommunity theory emphasizes the processes of dispersal between and persistence within patches (Leibold et al. 2004, Leibold and Chase 2017). These same themes are developed within parasite ecology, using the terminology (i) host-encounter filters and (ii) host-compatibility filters (Poulin 1997). From the parasite point of view, individual hosts are transient habitat patches that contain a community of parasites (an ‘infracommunity’ per parasite ecology (Bush and Holmes 1986, Poulin 1996)). The host population thus contains a single parasite metacommunity (a ‘component community’ in parasite ecology) that persists because parasites transmit from infected to uninfected individuals. This small-scale metacommunity is often nested within a larger metacommunity which is an assemblage of many distinct host populations. At either spatial scale, parasites must disperse between patches (host individuals or populations) and persist within those patches (Seabloom et al. 2015).

The processes that generate dispersal and persistence filters should differ by spatial scale in a parasite metacommunity (Fig. 1). For reasons elaborated below, some filters will act both among host individuals and among host populations, while others might act only at one scale. Such scale-dependent assembly rules might cause a parasite metacommunity to be more Clementsian at one spatial scale and more Gleasonian at another. We therefore tested whether patterns of co-occurrence among parasite species and the effects of biotic and abiotic variables on species distribution are scale dependent. Although there are many published studies of parasite metacommunities and of abiotic and biotic regulation of parasite distributions (Ebert et al. 2001, Mihaljevic 2012, Raeymaekers et al. 2013, Richgels et al. 2013, Dallas and Presley 2014, Seabloom et al. 2015, Borer et al. 2016, Cirtwill et al. 2016, Johnson et al. 2016, Hayward et al. 2017), to our knowledge this is one of the few and most extensive studies of how spatial scale alters the assembly rules of a spatially nested parasite metacommunity.

**Figure 1.**
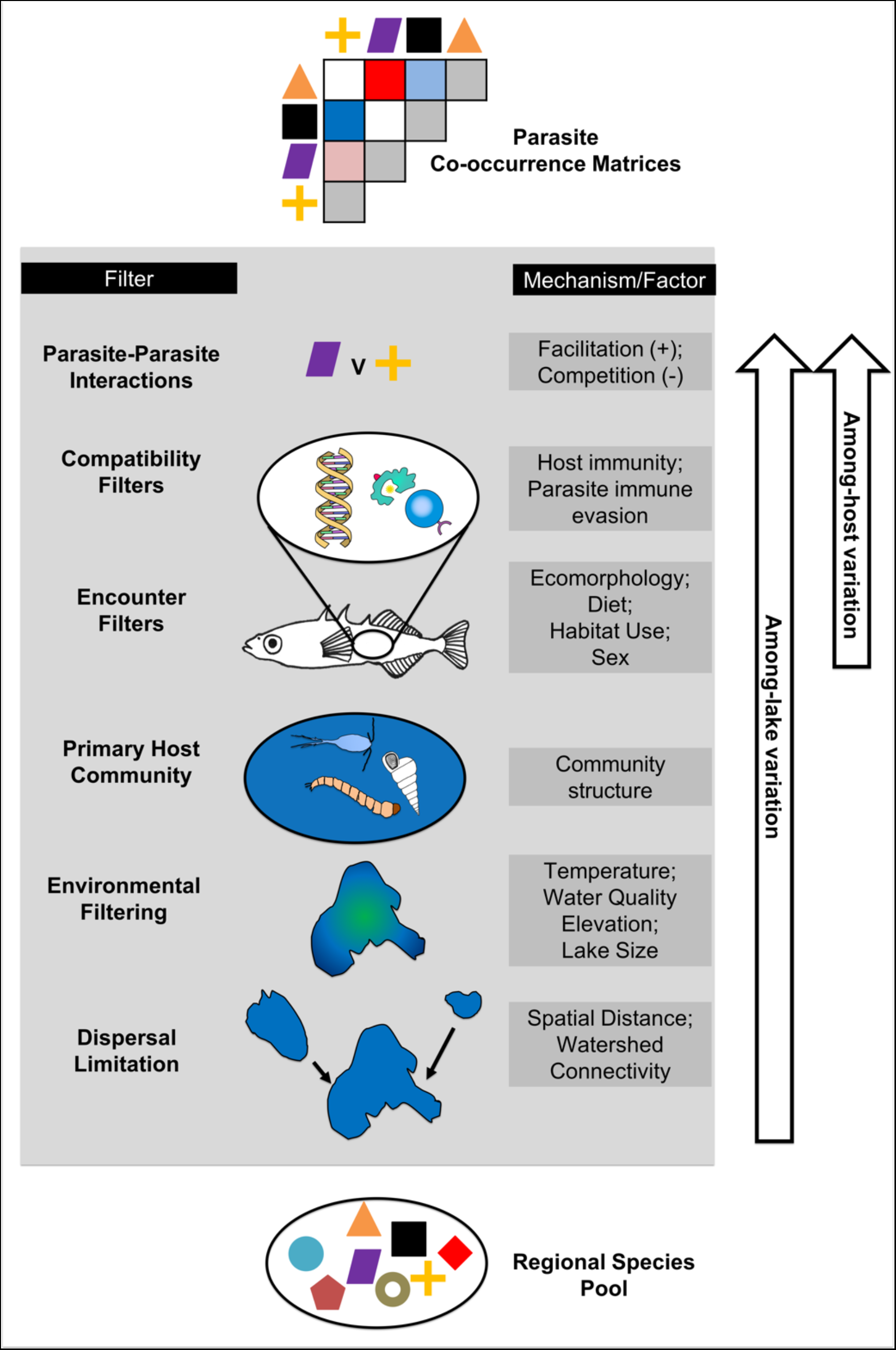
A schematic of factors contributing to parasite species’ distribution, co-occurrence, and metacommunity composition. We list examples of host individual and host population traits (right column) underlying general filters (left column). Arrows indicate the spatial scale at which the factors listed in the diagram are likely to act.

### Scale-independent factors

Some dispersal and persistence filters should act similarly across scales. Consider parasites with complex life cycles, where a focal host species must ingest an infected prey to be exposed to a parasite. Host populations often show ecomorphological and behavioral variation between individuals, leading to within-population variation in diet (Bolnick et al 2003). This ecological variation can cause host individuals to encounter different parasites at different rates (Hausfater and Meade 1982, Lafferty 1992, Wilson et al. 1996, Hutchings et al. 2003, Skartstein et al. 2005, Johnson et al. 2009, MacColl 2009, Johnson and Thieltges 2010, Stutz et al. 2014). For example, limnetic-feeding stickleback eating planktonic copepods are more likely to ingest copepod-transmitted *Schistocephalus solidus* cestodes, compared to benthivorous individuals in the same population (Stutz et al. 2014). At the host population scale, mean diet should predict exposure to and infection by diet-transmitted parasites. Thus, we expect ecologically-driven dispersal filters to have similar effects across scales.

### Among-host-population factors

A unique and fascinating feature of parasite meta-communities is that the host “patch” actively adjusts to kill or expel colonizing parasites. The capacity to resist parasites can be acquired by individuals after initial exposures (the adaptive immune response). Populations also evolve resistance to their most severe or commonly-encountered parasites, generating among-host-population variation in resistance, which in turn alters the prevalence of successful parasites (Maizels and Yazdanbakhsh 2003, Viney et al. 2005, Schmid-Hempel 2009). Because populations evolve, but individuals do not, evolution generates a uniquely population-scale source of variation in encounter filters (avoidance behaviors) or compatibility filters (host immunity) for most or all hosts in a population (Best et al. 2009, Berenos et al. 2010, Gilman et al. 2012, Luijckx et al. 2013). Thus, host or parasite local adaptation will tend to generate heritable differences in parasite communities (Hoeksema and Forde 2008) but only at the among-population spatial scale. This evolved immunity can undermine the relationship between encounter and infection rates. If hosts evolve effective resistance to their most frequently-encountered parasites, realized infection rates for a given parasite can actually be lowest where the parasite is most common. Herd immunity generates another uniquely population-scale effect. When a large enough fraction of hosts in a population is immune to a given parasite, even susceptible individuals are protected because their exposure rate declines.

Abiotic conditions represent another source of among-population variation in parasite community structure (Ebert et al. 2001, Richgels et al. 2013, Dallas and Presley 2014, Cirtwill et al. 2016). With a population, most individual hosts experience approximately similar abiotic conditions, so abiotic effects should be primarily a large-scale consideration. Likewise, ecological communities can differ across a landscape, changing parasite infection rates via the abundance of suitable and unsuitable hosts (dilution effect; (Johnson and Thieltges 2010, Becker et al. 2014)).

### Within-host-population factors

Filters might also generate metacommunity structure only at small scales (among individual hosts within a population). Sexual dimorphism in infection is common in natural populations (e.g., Reimchen and Nosil 2001)). Males and females systematically differ in diet (Shine 1989) and immunity (Rolff et al. 2009), which should contribute to within-population variation in infection. Assuming most host populations have an equal sex ratio, parasite metacommunity structure due to sexual dimorphism should be restricted to within-population scales.

To summarize, abiotic variables, community features, and host traits should structure parasite metacommunities. Some factors should act primarily at the among-host-individual scale (sexual dimorphism) and others at the among-population scale (evolved or herd immunity, abiotic conditions, dilution effects). Some factors should act similarly across spatial scales (diet increasing infection risk from prey-transmitted parasites). These scale-dependent effects should then dictate which parasite species tend to co-occur, or not.

To test these expectations, we document the metacommunity structure of a multi-species parasite assemblage infecting threespine stickleback (*Gasterosteus aculeatus*). This is an appealing study system because stickleback populations (e.g. in separate lakes) differ non-randomly in parasite community diversity (Eizaguirre et al. 2011), composition (Poulin et al. 2011, Stutz et al. 2015), and infection intensity (Pennycuick 1971, Weber et al. 2017). Differences in infection among stickleback populations have been shown to be temporally stable (Weber et al. 2017, Young and MacColl 2017), and have been linked to differences between stickleback populations in immune genotype (Matthews et al. 2010, Eizaguirre et al. 2012b, Stutz and Bolnick 2017), diet (Matthews et al. 2010), and abiotic conditions (e.g., salinity; Simmonds and Barber 2015). Within populations, individual stickleback infection is correlated with individual diet, ecomorphology, sex, and immune genotype (Reimchen 1997, Reimchen and Nosil 2001, Wegner et al. 2003, Matthews et al. 2010, Eizaguirre et al. 2012a, Stutz et al. 2014, Stutz and Bolnick 2017). However, these prior studies have tended to focus on one or a few variables at a time, and are generally restricted to a single spatial scale. Perhaps because of this scale-limited scope, there have been conflicting conclusions among the studies cited above.

Here, we identify abiotic, genetic, phenotypic, and ecological features of host individuals and host populations that help explain among-individual and among-population variation in parasite composition. We identify scale-independent and scale-dependent factors influencing parasite community composition. A related paper (Bolnick et al. Manuscript) considers scale-dependence of factors regulating parasite species richness, an emergent property of the processes considered here.

## Methods

### Collection

In June 2009, we collected stickleback from 33 lakes in 9 watersheds on Vancouver Island, British Columbia, Canada (details in Table S1, Figure S1), in the historical lands of the Kwakwaka’wakw First Nations. Collection and animal handling were approved by U.T. IACUC (#07-032201) and a Scientific Fish Collection Permit from the Ministry of the Environment of British Columbia (NA07-32612). We also sampled from 8 streams and 5 anadromous populations in estuaries, but for most analyses here we focus on lake population data. Sites were chosen non-randomly to sample a broad array of lake types within a small geographic region. We placed unbaited 0.5-cm gauge wire minnow traps along ∼200 m of shoreline in 0.5-3m deep water. We obtained 60-100 fish per site (Table S1). Fish were immediately euthanized in MS-222 and preserved in 10% buffered formalin after cutting a fin clip into ethanol for DNA. Specimens were rinsed and stored in 70% isopropyl alcohol after staining with Alizarin Red.

### Data acquisition 1: parasite infections

We counted macroparasites (helminths, crustaceans, molluscs, and microsporida) in each fish with a stereodissection microscope. We scanned the skin, fins, and armor plates, and then the buccal cavity and gills. We then dissected the body cavity and organs (liver, swim bladder, gonads, eyes) and opened the digestive tract. Parasites were identified to the lowest feasible taxonomic unit (typically genus). For abundant gill parasites, we counted parasites only on the right side. For each taxon, we calculated per-population infection prevalence (proportion of fish infected) and abundance (mean number of parasites per fish) following (Bush et al. 1997), and confidence intervals of proportions following Newcombe (1998).

### Data acquisition 2: stickleback morphology

We quantified stickleback ecomorphology, which is known to covary with individual diet within populations (Robinson 2000, Snowberg et al. 2015) and among populations (Lavin and McPhail 1985, Lavin and McPhail 1986). Before necropsy, we weighed all fish to 0.01g and used digital calipers to measure external body dimensions (in mm): standard length, body depth, and body width at the pectoral fins. For a subset of ∼30 individuals per population, we measured trophically important traits: gape width, gill raker number, and length of the longest gill raker. We inspected gonads via dissection to determine sex. Linear measurement data were log transformed and size-standardized by regression on log standard length.

### Data acquisition 3: stickleback diet

For a random subset of 28 populations, we analyzed stickleback stomach contents for recent diet. Previous studies have shown that individual sticklebacks’ stomach contents are indicative of long-term diet as inferred from stable isotopes, morphology, and feeding observations in the wild (Snowberg et al. 2015). We removed stomachs from the same fish measured for morphology, and identified the presence/absence of each prey taxon to the lowest feasible taxonomic level (typically family). For analysis, we binned prey taxa into functional groups (benthic or limnetic) and calculated the proportion of benthic prey in each fish’s stomach. For each population, we calculated the average proportion of benthic prey across the sampled individuals.

### Data acquisition 4: stickleback genetic diversity

To quantify the effect of host genetic variation on parasite distributions, we used ddRADseq (Peterson et al. 2012) to obtain single nucleotide polymorphism (SNP) genotypes from a subsample of 12 fish from each of 31 lakes (Table S1 in Bolnick et al, manuscript). Protocols, bioinformatics steps, and SNP filtering are exactly as described in Stuart et al. (2017). The result was a matrix of genotype scores for 175,350 SNPs in 336 fish (averaging 107,698 SNPs per individual). (36 individuals were dropped due to poor sequence coverage.) We calculated genome-wide heterozygosity for each fish, and between-population genetic distances (Weir-Cockerham unbiased F_ST_).

## Data Analysis

As described in detail below, we began by testing for non-random co-occurrence between parasite species. We considered co-occurrence at the level of host individuals within populations, then among host populations, then tested whether co-occurrence is similar across these scales. Next, we tested for individual host or host population characteristics that might act as dispersal or persistence filters that affect parasite species distributions. Last, we tested whether different parasite species are subject to similar filters, explaining their co-occurrence or lack thereof.

### Analysis 1: Do certain pairs of parasite species tend to co-occur within hosts?

We estimated a co-occurrence matrix between parasite species within each of the 33 host populations. This matrix measures the tendency for pairs of parasite species to infect the same host individuals (Fenton et al. 2010). We calculated Spearman rank correlations between all pairs of common parasite taxa (i.e. infecting >5 fish within a population). The Holm-adjusted p-values from these rank correlations test the null hypothesis that the pairwise combinations of parasite species are independently distributed among individual hosts within a given host population.

Patterns of co-occurrence between parasite taxa might be inconsistent between host populations. For each possible pairwise comparison of populations, we calculated a Mantel correlation between their respective co-occurrence matrices. A significant positive correlation implies that similar parasite combinations co-occur in both host populations (Poulin 2007, Presley 2011, Meynard et al. 2013). We used only parasite taxa found in both populations.

### Analysis 2: Do parasite species tend to co-occur within host populations?

If patterns of co-occurrence are scale-dependent, parasite co-occurrence matrices should differ within versus between host populations. To test this, we calculated the mean parasite abundance (*sensu* Bush et al (1997)) for each taxon in each lake. We then calculated the Spearman correlation between the mean abundances of each pair of parasite taxa, at the host population level. Each pairwise comparison was tested against a null hypothesis of independence.

Next, we tested whether within-population co-occurrence and between-population co-occurrence matrices are similar. For each host population, we used a Mantel test to compare the focal population’s individual-level co-occurrence matrix (estimated in Analysis 1) versus the between-population co-occurrence matrix (preceding paragraph). Low correlations suggest that the processes generating co-occurrences are scale-dependent.

Note that because co-occurrence patterns were inconsistent among populations, and across spatial scales (see Results), we did not use community ordination (e.g., RDA or NMDS) to summarize multivariate community variation among individuals or among populations.

### Analysis 3: What traits of host individuals predict within-fish parasite community structure?

We used negative binomial mixed model GLMMs to relate each parasite taxon’s abundance as a function of host traits, with host population as a random intercept and random slope. For the full dataset of all hosts, we tested for sex and mass effects. For the 30 individuals per population with detailed morphological data we tested for effects of sex, mass, gill raker number, gill raker length, and gape width. For the subset of populations with diet data, we ran similar GLMMs adding host diet Principal Component axis 1 and 2 as a model predictor, again with host population providing a random intercept and slope. Sample sizes for these models are given in Supplementary Table S1.

Last, we tested for individual host genotype effects on parasite prevalence using a genome-wide association study (GWAS). For this GWAS, for each SNP we used a binomial GLM testing whether the presence of infection by a given parasite depended on individuals’ genotypes at that SNP, with host population as a fixed effect to control for among-population covariation in infection and genotype. Using false discovery rate correction, we iterated this analysis across all SNPs that were scored on at least 25% of the sampled fish and had a region-wide minor allele frequency exceeding 5% (for populations where the focal SNP was polymorphic). We only analyzed parasites found in at least 5 populations.

### Analysis 4: What features of host populations predict across-site parasite community structure?

We used variation-partitioning (Borcard and Legendre 2002, Cottenie 2005, Peres-Neto et al. 2006, Logue et al. 2011) to find abiotic and biotic factors that contribute to among-host-population variation in parasite metacommunity composition and estimate how much variation in multi-species community composition can be partitioned to among-host-population spatial distance, genetic distance (F_ST_), ecomorphology, and environment (for this one analysis we included stream fish). We excluded two lakes and one stream for which we did not obtain sufficient sequence data (Browns Bay Lake, Farewell Lake, Farewell Stream). We also excluded two sites from a separate island (Quadra – Village Bay Lake and Village Bay Stream), whose geographic distance had high leverage on the distance effect estimate, leaving a total of 36 sites. We were interested in two spatial distance matrices: a ‘fish swims’ spatial distance calculated along tributaries that connect two sites, and over-land Euclidean distance. For spatial and genetic data, distance matrices were converted into rectangular data through Principal Coordinates of Neighborhood Matrix (PCNM), which computes a principal coordinate analysis using a truncated distance matrix. For PCNM, we used the function pcnm() in the package vegan and extracted eigenvectors with positive Moran’s index of autocorrelation (Dray et al. 2006). We then ran redundancy analysis (RDA) on each component (spatial, genetic, ecomorphology, environment) to determine if that component significantly explained Hellinger transformed parasite community data. Over-land euclidean distance was not significant and not included in variation partitioning analyses. For all other components, we ran forward selection on that component to determine which variables to include. We retained log gape width residuals and condition for ecomorphology and retained habitat and maximum depth for environment. We then ran the function varpart() in the package vegan to partition variation in community data between these four components.

Next, we examined each parasite taxon separately to identify lake-level abiotic and host phenotypic traits associated with each parasite taxon’s prevalence. Using lake as the level of replication, we used binomial GLMs to regress parasite prevalence against site characteristics (lake area and elevation) and host population characteristics (means of fish mass, gill raker length, and gape width). For the subset of lakes with diet data, we re-ran these analyses, adding the top two axes of diet variation as independent variables.

To evaluate genetic contributions to among-population variation in infection rates, for each SNP we used a binomial GLM to test whether each parasite taxon’s prevalence was a function of that SNP’s allele frequency (using population as the level of replication), with watershed as a covariate. We calculated q-values to adjust for multiple comparisons.

### Analysis 5. Do host population traits explain patterns of parasite co-occurrence ?

We hypothesized that similarities in host encounter or compatibility filters, across parasite taxa, are associated with higher parasite co-occurrence at the host population scale. For instance, parasites that respond similarly to lake size will tend to co-occur. To test this, we took the effect size estimates (Z scores) from the GLMs in Analysis 4, which represent the effect of lake characteristics and host population trait means on each parasite taxon. We then calculated the correlation between these effect sizes for each pair of parasite taxa. These correlations generated a ‘co-dependence’ matrix expressing the similarity, between each pair of parasite taxa, in their dependence on host and environmental traits (at the host population scale). We then used a Mantel test to evaluate the correlation between this co-dependence matrix and the across-host-population co-occurrence matrix (Analysis 2). We restricted this analysis to the among-population scale because within population co-occurrence patterns were not repeatable among host populations.

## Results

We observed striking variation in parasite infections among stickleback populations and among individual fish within populations (Fig. 2). For example, Unionidae range from as low as 0% prevalence (e.g., in Higgens lake, CI 0%-5%), to 100% prevalence (e.g., in Little Mud Lake, CI 93%-100%). Within a given lake, some individuals had zero Unionidae while other fish had up to 120 covering their gills. Although only a few taxa spanned such a wide range, every parasite had highly significant among-population differences in prevalence (binomial GLMs, all P < 0.0001).

**Figure 2.**
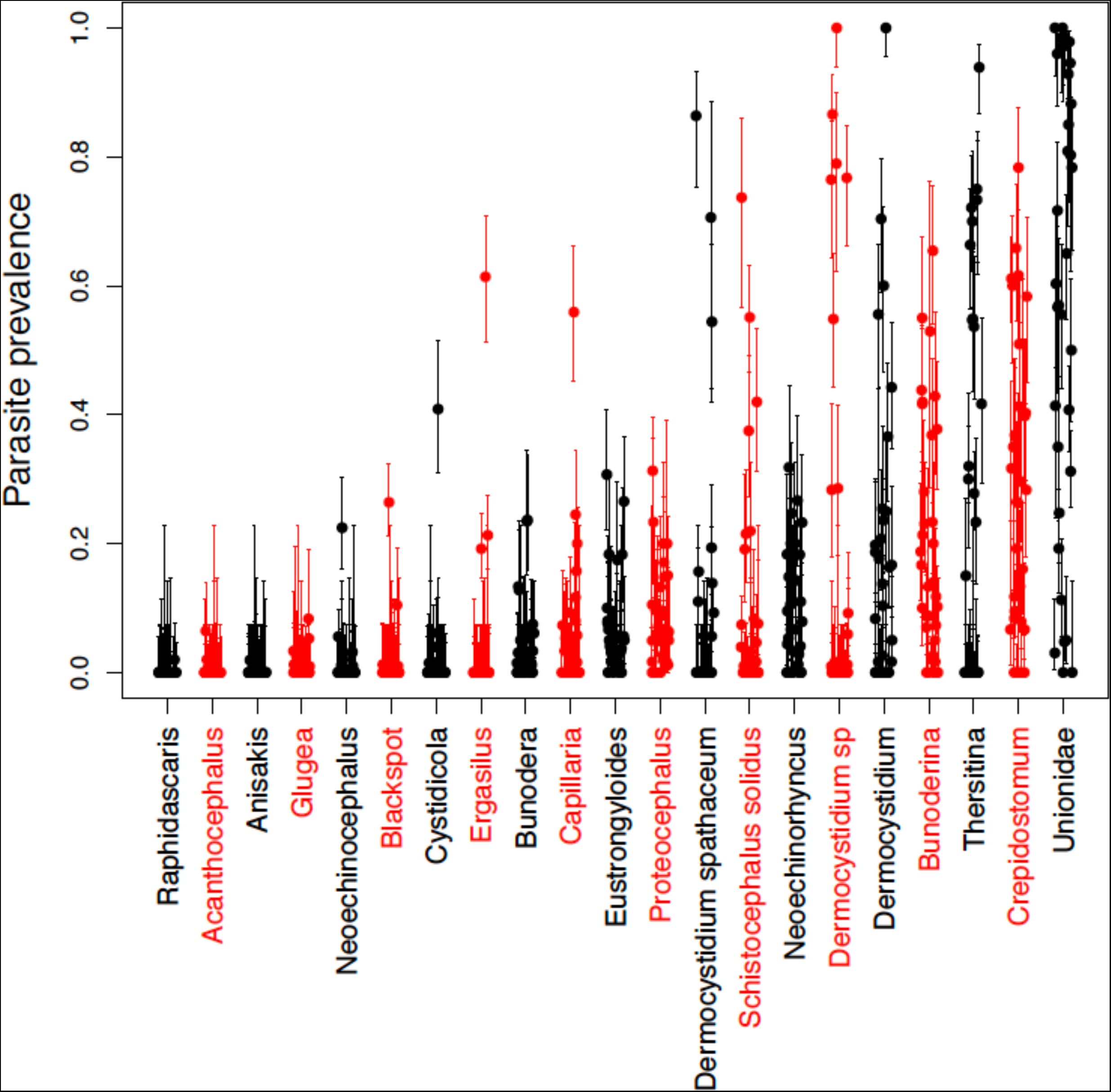
Variation in parasite prevalence among 33 lake populations of threespine stickleback. Each point represents the proportion of fish infected by the focal parasite, with standard error bars. Parasites are ordered along the x axis from least to most common in the metacommunity.

### Analysis 1: Parasites co-occur within hosts, but inconsistently across host populations

We found evidence for significant parasite co-occurrence across fish, but these co-occurrence patterns did not repeat themselves across lakes. For instance, in McCreight Lake, 8 out of 91 pairwise comparisons between parasite taxa were significantly positively correlated (Fig. 3A), such as the abundance of *Thersitina* and Unionidae (both horizontally-transmitted external parasites on gills; Fig. 3B). Although most populations had multiple significant, positive pairwise correlations (median = 8 different parasite pairs, Fig. S3), there was wide variation between populations. For example, we found 41 significant parasite-parasite correlations out of 231 pairwise comparisons in Roberts Lake (∼70% positive; Fig. S2), whereas we observed only eight parasite taxa and no pairwise correlations in Cecil Lake (200m upstream from Roberts Lake).

**Figure 3.**
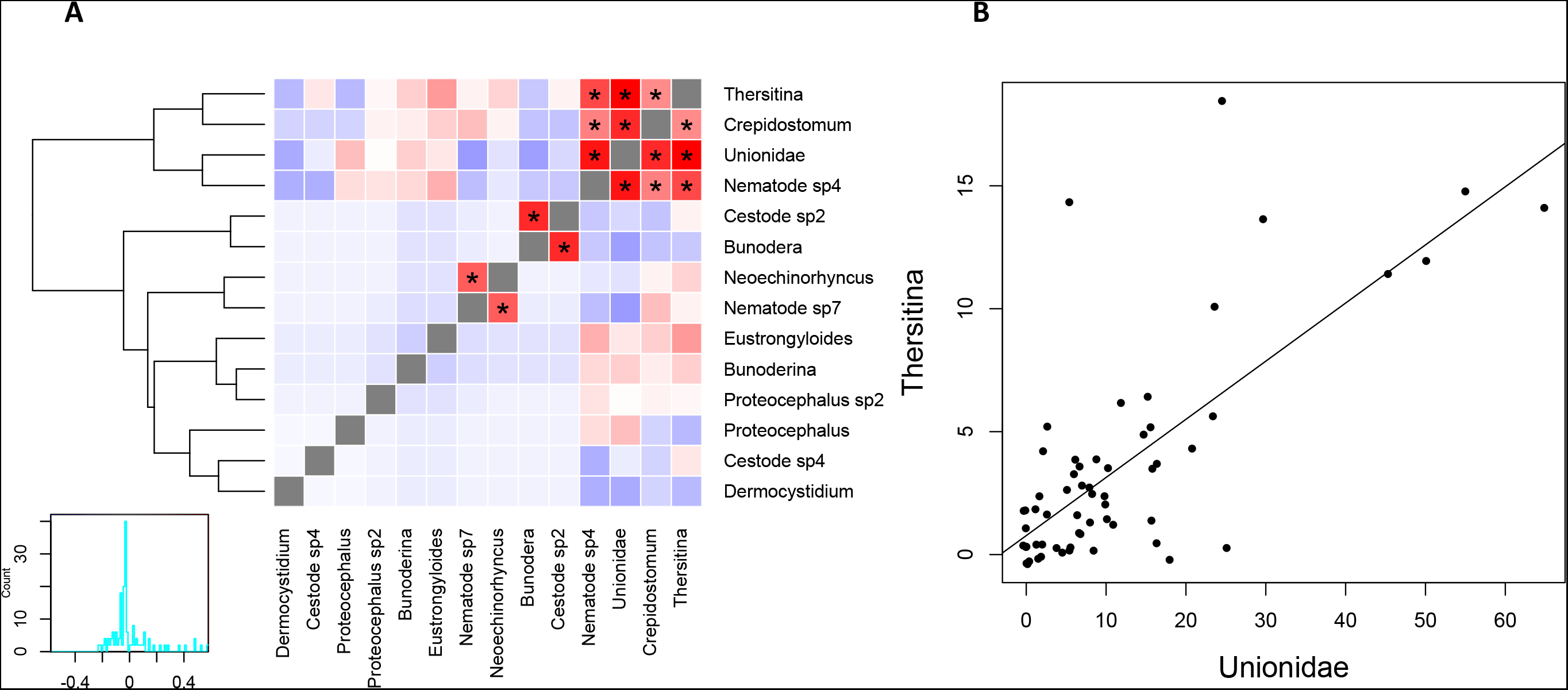
(A) The parasite species co-occurrence network observed among individual stickleback within McCreight Lake (N = 60 fish). Red and blue shaded cells show positive and negative covariance, respectively; asterisks denote significant Spearman correlations. A dendrogram clusters species with more similar co-occurrence. (B) A scatter plot of the correlation between the abundance of *Thersitina* and Unionidae infections. Points represent host individuals. There are overlapping points at (0,0).

The strength and identity of parasite co-occurrence differed among stickleback populations (Fig. S3). The average Mantel correlation between two lakes’ co-occurrence matrices was only r_M_=0.028. Of 561 possible between-population comparisons, 17% exhibited significant positive correlations (6% were significantly negative). This weak generality reflects many cases where two parasites were correlated in some lakes, but not others even when they were both present. For example, *Dermocystidium* and *Thersitina* were positively correlated in four lakes (three different watersheds) but not in 10 other lakes where they were both present (Fig. S3). Sometimes parasites were positively correlated in some lakes and negatively in others (e.g. *Thersitina* and Unionidae), indicating that processes driving co-occurrence at the host-individual scale are inconsistent across the metacommunity.

### Analysis 2: Parasites tend to co-occur within host populations

We found strong co-occurrence of parasite mean abundance at the scale of host populations (Fig. 4). For example, *Crepidostomum* (an internal helminth with a complex multi-host life cycle) and Unionidae (a directly transmitted mollusk gill parasite) tend to be either both common or both relatively rare in populations (Fig. S4). Notably, among-lake co-occurrence involves stronger and more significant correlations than we observed within any single lake.

**Figure 4.**
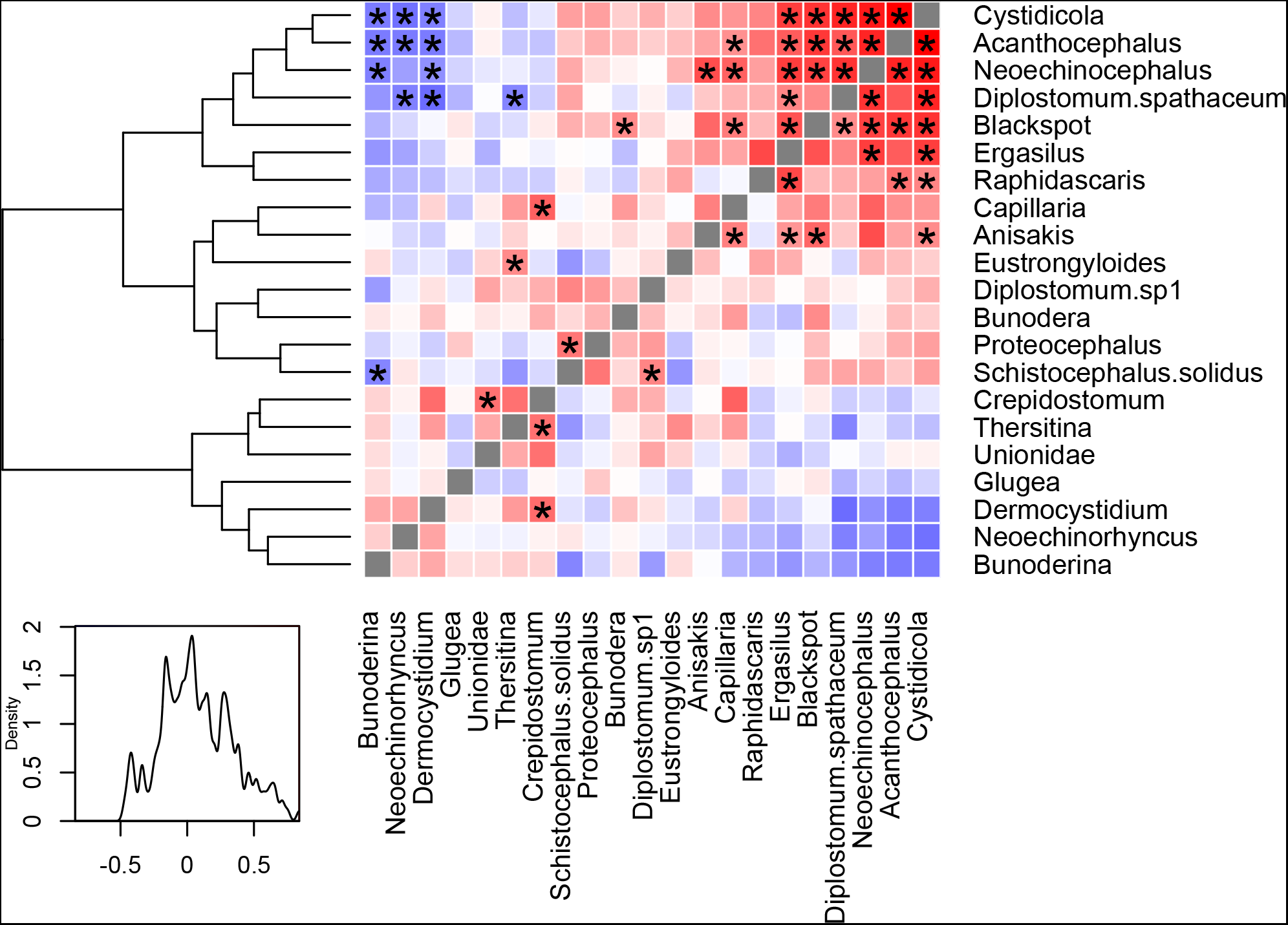
Parasite species co-occurrence across host populations. Red denotes positive, and blue negative, Spearman rank correlations between parasite taxa, using host population (lake) as the level of replication (scale with density trace is provided, bottom left). The dendrogram indicates sets of parasites that covary positively. Asterisks indicate statistically significant correlations (P < 0.05 above diagonal, Bonferroni-corrected below diagonal). An example of parasite co-occurrence among lakes is plotted in Fig. S4.

The among-lake co-occurrence matrix was not related to the within-lake co-occurrence matrices. The Mantel correlations between these spatial scales’ matrices ranged from −0.39 to 0.34 (depending on the focal lake for individual-scale co-occurrence), and on average were indistinguishable from zero (mean = −0.046, sd = 0.25). These results imply that among-lake covariation in parasite community structure is a result of non-random community assembly processes at both scales, but these processes differ between the individual host, and host population scales.

### Analysis 3: Individual host traits predict parasite community structure

Ecomorphological characteristics explained variation in parasite abundance among individual hosts for many parasite taxa (see Table S2 for statistical details). The strongest trend was for parasites to be more abundant in larger fish (e.g., Fig. S5A), as has been found in other host species (Timi and Poulin 2003). This trend might reflect age, higher feeding rates, or a particular diet (larger stickleback eat more benthic prey). Associations between parasites and other trophic traits suggest that fish diet affects individual parasite loads: individuals with larger gapes (a benthic trait) had more abundant *Capillaria* and *Crepidostomum*, but fewer *Ergasilus*, *Neoechinorhyncus* and *Schistocephalus*. More numerous gill rakers (a limnetic trait) coincided with more *Crepidostomum* but fewer *Ergasilus*. Longer gill rakers (also a limnetic trait) conferred more *Blackspot*, *Bunoderina*, *Proteocephalus*, and *Schistocephalus*, but fewer *Ergasilus*, *Eustrongylides*, and Unionidae (Fig. S5B).

Stomach contents were also associated with infection. Individual stickleback with more limnetic diets (higher diet PC1 scores) had more *Schistocephalus* (Fig. S5C), but fewer *Eustrongylides*, consistent with these parasites’ limnetic and benthic first hosts and corroborating a prior study (Stutz et al. 2014). Host sex affected infection rates for several parasites (Table S2), typically with higher infection rates in females, who also had higher parasite richness (Bolnick et al manuscript). *Schistocephalus* was the sole species that was significantly more common in males than in females, consistent with males’ tendency towards a more limnetic diet (Reimchen and Nosil 2001, Snowberg et al. 2015). Our individual-level GWAS analysis found no SNPs that correlated significantly with individual-level infection after controlling for population-level variation in both allele frequency and infection. GWAS will fail to identify loci when its assumption of parallel evolution is not met.

### Analysis 4: Host population traits predict parasite community structure

Variance partitioning analysis revealed that spatial distance, environment, genetic distance, and host ecomorphology each explained a significant portion (p < 0.05) of among-population variation in parasite community composition. Host ecomorphology explained the most variation, and spatial distances the least (Fig. S6).

For individual parasite taxa, lake biogeography strongly affected the distribution of multiple parasite species across populations (Table S3). Larger lakes had fewer Unionidae, *Crepidostomum* (Fig. S7A), and *Dermocystidium*. Higher-elevation lakes had higher prevalence of *Dermocystidium*, *Schistocephalus* (Fig. S7B), *Diplostomum*, and *Cystidicola*, and lower prevalence of *Ergasilus* (Table S2). Lakes farther from the ocean had more *Crepidostomum*, *Dermocystidium*, and *Ergasilus* but fewer *Diplostomum*. More remote lakes (farther from the ocean) tended to have higher infection by most parasites (mean Z score across all parasites = 1.95, P = 0.066). These effects are summarized with a heatmap in Fig. 5A.

**Figure 5.**
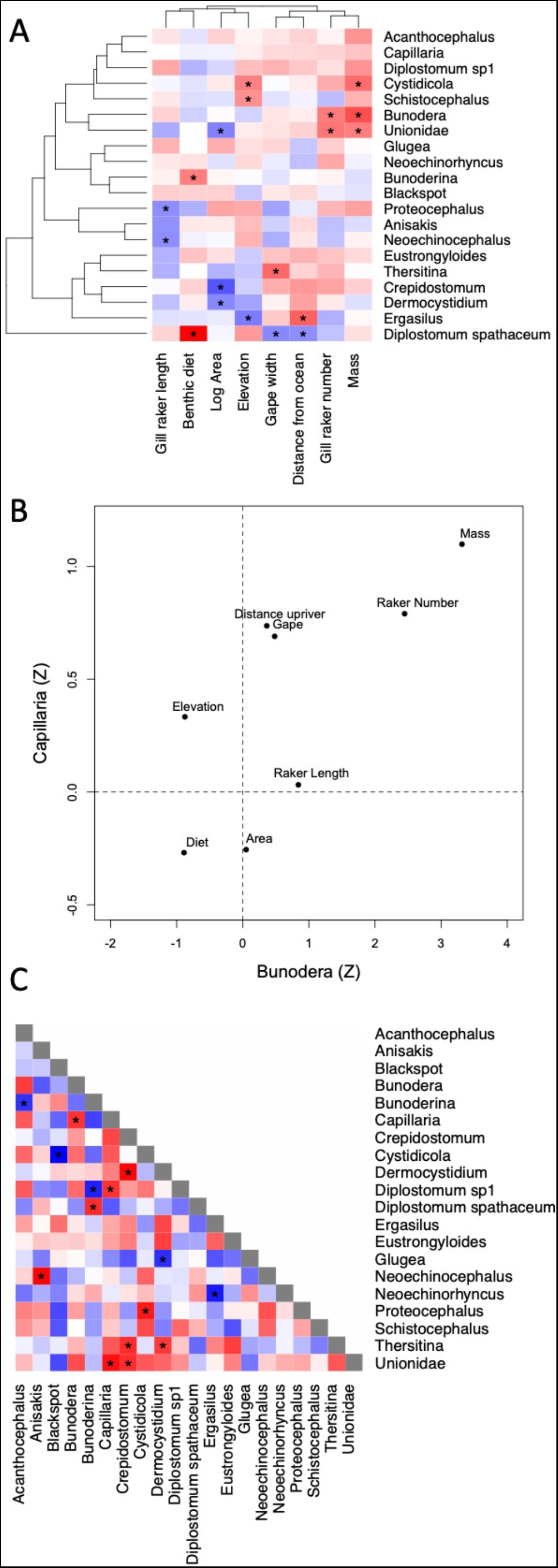
A) Heatmap of parasite dependence on host population characteristics, for the most common parasites. Red denote positive correlations, blue negative. Asterisks show significant associations. B) The correlated effects (Z statistics from the GLM) of host population characteristics on two parasite taxa, *Bunodera* and *Capillaria*. C) Co-dependence matrix—the correlation between parasite species’ associations with environmental and host population traits (e.g., summarizing all pairwise comparisons as in in panel B). We calculated correlation coefficients between Z-scores from each factor in the linear models for each species.

Populations with larger fish (higher mean mass) had more Unionidae (Fig. S7C), *Diplostomum*, *Proteocephalus*, *Bunodera*, *Cystidicola*, and *Acanthocephalus*. The positive effect of mean fish mass is consistent across most parasite taxa (mean Z = 3.12 P = 0.006). Only half of these associations were mirrored at the individual host level (Unionidae, *Diplostomum*, and *Proteocephalus* were positively related to individual fish mass). Fish diet and ecomorphology also influenced infection at the among-population-level: more limnetic populations (higher diet PC1) were more heavily infected by *Bunoderina* (Fig. S7D) and *Diplostomum spathaceum* (an association not seen at the individual host scale). *Proteocephalus* was associated with gill raker length at both spatial scales, but the direction of the effect changed from positive to negative with increasing scale. Populations that on average had more gill rakers (a typical limnetic trait) had more Unionidae (Fig. S7E) and *Bunodera*, neither of which were associated with gill raker number at the individual host scale. Populations with larger gape widths had more *Thersitina* (Fig. S7F) but fewer *Diplostomum*; neither associated with gape at the individual host scale.

GWAS at the among-population scale revealed numerous significant correlations between population allele frequencies at a given SNP and parasite prevalence. We ran 1,281,483 GLM tests (for all combinations of sufficiently polymorphic SNPs and common parasites). Applying a significance threshold of *α* = 5*10-14 (more stringent than Bonferroni), we located 14,832 SNP-parasite associations, examples plotted in Fig. S8. These SNPS correlated disproportionately with a few parasites (max = 1085 SNPs), while other parasites had no significant associations. This result indicates that host genetic variation is associated with parasite infection across populations.

### Analysis 5. Co-dependence on population-level traits explains patterns of parasite co-occurrence

Some groups of parasite taxa depended on the same sets of host-population traits in Analysis 4 (Fig. 5A). For example, *Capillaria* and *Bunodera* were both more common in lakes where fish were larger and had more gill rakers, and less common in lakes with a more limnetic diet. The effects of host population traits on these two parasites were highly (r = 0.722, P = 0.042, Fig 5B). Such ‘co-dependence’ indicates similar effects of host population traits (Fig. 5C). Conversely, *Cystidicola* and Blackspot had negative co-dependence, responding to similar host traits but in opposite directions (r = −0.913, P = 0.002, Fig. S9).

Parasite taxa that showed positive co-dependence on host population traits were more likely to co-occur at the scale of host populations. That is, there was a significant positive correlation between co-occurrence (Fig. 4) and co-dependence (Fig. 5C), as illustrated in Fig. 6 (Mantel r = 0.34 P = 0.001). This result implies that the non-random structure in the higher-scale parasite metacommunity is generated by shared parasite responses to host population characteristics.

**Figure 6.**
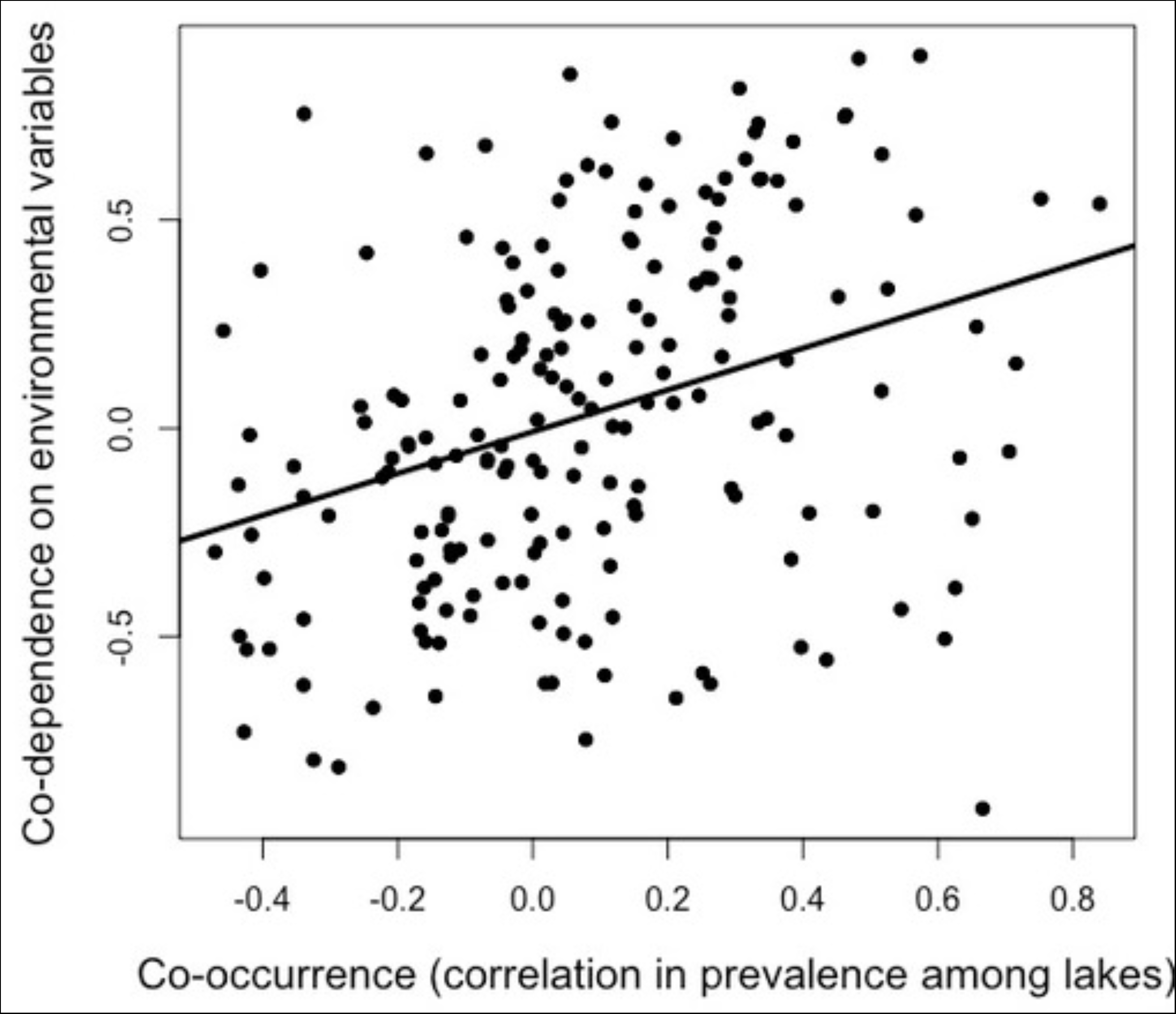
Correlation between the co-occurrence matrix (Fig. 4) and the co-dependence matrix (Fig. 5C). The line represents a significant linear model fit (Mantel test; P = 0.001).

## Discussion

Parasites form hierarchically structured metacommunities, whose composition varies among host individuals within populations, and between host populations (Poulin 2007, Mihaljevic 2012, Seabloom et al. 2015, Borer et al. 2016). Here, we have shown that lake characteristics, and host ecology and genotype, jointly structure a stickleback parasite metacommunity. The particular factors affecting parasite species’ abundance are scale-dependent, however.

The stickleback parasite metacommunity examined here is moderately Clementsian: some species tend to co-occur more often than expected by chance. This co-occurrence is most pronounced at the among-host-population scale. At the smaller spatial scale, certain parasite species do tend to co-occur, but such pairs are rarer, weaker, and tend to be positively correlated. The implication is that, as one moves to the larger spatial scale, the parasite community becomes more deterministically structured by habitat patch characteristics (lake features or host population trait means). Consistent with this inference, parasite variation among host populations is more strongly regulated, by a greater number of biotic and abiotic factors, than variation among host individuals.

### Spatial scale and the factors affecting parasite community structure

Parasite metacommunity structure can broadly be explained by a combination of dispersal and persistence filters, equivalent to what parasitologists call encounter and compatibility filters, respectively (Fig. 1). We found that parasite communities depended on two interrelated ecological encounter filters: host diet and trophic morphology. In a few cases these ecological filters acted at both spatial scales. For example, *Diplostomum* was more common in larger host individuals and in larger (on average) host populations, a trend found in 3 of the 21 parasite taxa examined. This trend is consistent with body size effects seen in many other fish species (Timi and Poulin 2003), perhaps reflecting greater age and greater time to accrue parasites. *Proteocephalus* also shared an ecological filter (gill raker length) at both spatial scales, but the sign of this correlation was reversed. More typically, however, host diet and morphology had effects on parasite abundance at one scale, but not another. This scale-dependence is counter to our expectation that trophically-transmitted parasites should exhibit similar ecological encounter filters at both spatial scales. This surprising result might be due to scale-specific effects of other filters (e.g., immunity) that counter-act the ecological filters at one scale but not another.

Some filter mechanisms can only be significant at one spatial scale and thus could cause scale dependence. For instance, males and females differed in parasite infections, consistent with previous studies in stickleback (Reimchen and Nosil 2001). Because our populations do not differ greatly in sex ratio, sexual dimorphism should primarily contribute to within population metacommunity variation. Conversely, geographical characteristics of entire lakes (size, elevation, distasnce from ocean) are necessarily shared by all individuals within a given lake and so only contributed appreciable variation at the among-population scale.

In metacommunities, the geographic positions of habitat patches generate divergence in community structure by modifying the rates and sources of colonization. A previous study in Scottish lakes also identified between-population genetic dissimilarity as a major contributor to between-population differences in parasite community composition (Rahn et al. 2016). Our populations were spread over an ∼1,500 square km region (70 km north-south range, 45 km east-west range) and subdivided into watersheds. Our variance decomposition analysis found that distance via watersheds “as the fish swims” significantly explained parasite community composition but that Euclidean distance “as the crow flies” did not. However, neither effect was especially strong: population differences in host ecological traits and genetic divergence were appreciably more important.

### Coevolution

The variance partitioning analysis confirms that genetically divergent host populations tend to have more divergent parasite communities, controlling for the relatively weak confounding effect of spatial distance. This result suggests that there is genetic variation in infection risk, which arises in large part from shared ancestry (affecting whole-genome F_ST_), not just from selection on particular loci. But, evolved differences in host resistance (e.g., Weber et al. 2017a,b) can also contribute to parasite metacommunity structure. Host and parasite (co)evolution are most relevant to population-scale metacommunity structure, because natural selection acts on populations, not individuals. In fact, selection should have opposing effects on metacommunity variation at these scales. Selective sweeps within host populations simultaneously increase between-population differences and reduce within-population polymorphism (reducing among-individual genetic variation in resistance). Such sweeps will generate genotype-parasite associations at the among-population level, but remove variance to detect such effects within populations. Alternatively, parasites might impose balancing selection within host populations (e.g., Wegner et al. 2003), promoting among-individual associations between genotype and resistance, but inhibiting population divergence. Our genome-wide association study (GWAS) found numerous SNPs associated with infection variation among populations, but not among individuals within populations. This scale-dependent effect suggests that, in this system, parasites primarily drive divergent rather than balancing selection. We cannot at present determine whether this effect acts through immune genes, or loci affecting traits like size or diet.

Locally evolved host immunity could explain why dietary and trophic morphology effects are inconsistent across spatial scales. *Schistocephalus solidus*, for example, is contracted when fish eat infected copepods, so limnetic-feeding individuals are at greater infection risk (Stutz et al. 2014). The expected ecology-infection correlation holds at the host-individual scale (Table S2) but not the host-population scale (Table S3). Why are limnetic-feeding populations not more heavily infected? Perhaps, populations with high ecological risk of infection are more likely to evolve resistance to that parasite, which would dampen population-level differences in infection (Fleischer et al. Manuscript). There is clear evidence that certain lake populations of stickleback have evolved greater resistance to *S.solidus* (Weber et al 2017a,b). The individual-level effect persists within populations, where recombination breaks up any genetic associations between ecomorphology and immunity.

### Co-dependence and co-occurrence

The results described above indicate that most parasite taxa in this metacommunity are regulated by a combination of host traits at the individual-host scale, and at the between-host-population scale are regulated by abiotic conditions, host traits, and host allele frequencies. At the population scale, some combinations of parasites respond in the same directions to the same sets of population traits (e.g., show co-dependency). Other combinations of parasite taxa respond in opposite directions or are simply independent (Fig. 9). Parasites that are more co-dependent on the same host population characteristics are more likely to exhibit positive co-occurrence (e.g., be found in the same locations). We recognize an important caveat: co-dependence and co-occurrence estimates were derived from the same set of lakes. The sample sizes of the present study, while large, are not sufficient to warrant cross-validation between separate subsets of the data. A valuable goal in the future would be to revisit this pattern using separate training and test datasets to avoid potential for circularity.

### Summary

The stickleback parasite metacommunity studied here is structured by a wide variety of factors: lake geography, host mean traits, host allele frequencies, and individual traits (including sex). These factors are almost all scale-dependent (except host body size), which means that the mechanistic basis of infection and epidemiology cannot readily be generalized from individual animals to their populations, or vice versa. This provides a specific case study illustrating the potentially broader point that filters structuring metacommunities are highly scale-dependent. Such scale-dependent effects can explain inconsistent findings among studies conducted at eifferent scales, and imply that multi-scale studies should be the norm for parasite metacommunity ecology.

## Acknowledgements

We thank C. Harrison, T. Rodbumrung, and T. Ingram, for assistance with field work. J. Day assisted with parasite data collection. R. Grunberg provided valuable feedback on the manuscript. Research was supported by the David and Lucille Packard Foundation, a Howard Hughes Medical Institute Early Career Scientist award, NSF (DEB-1144773), and NIH (1R01AI123659-01A1) to DIB, and an NSF Graduate Research Fellowship to EJR.

---

**Figure S1.**
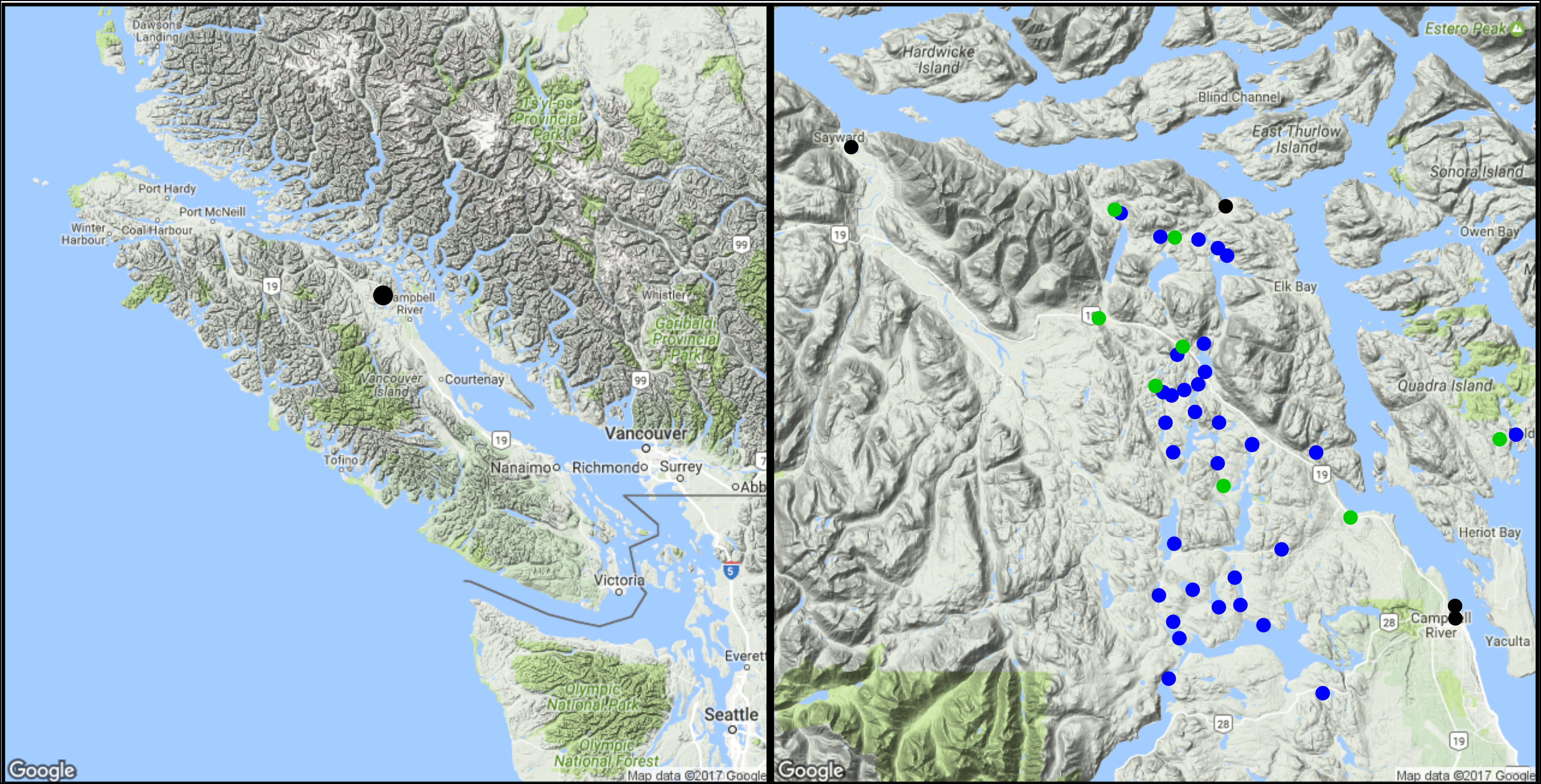
A) Vancouver Island, with a black dot indicating the location of the inset map (B), where lake, stream, and marine sample sites are indicated in blue, green, and black points.

**Figure S2.**
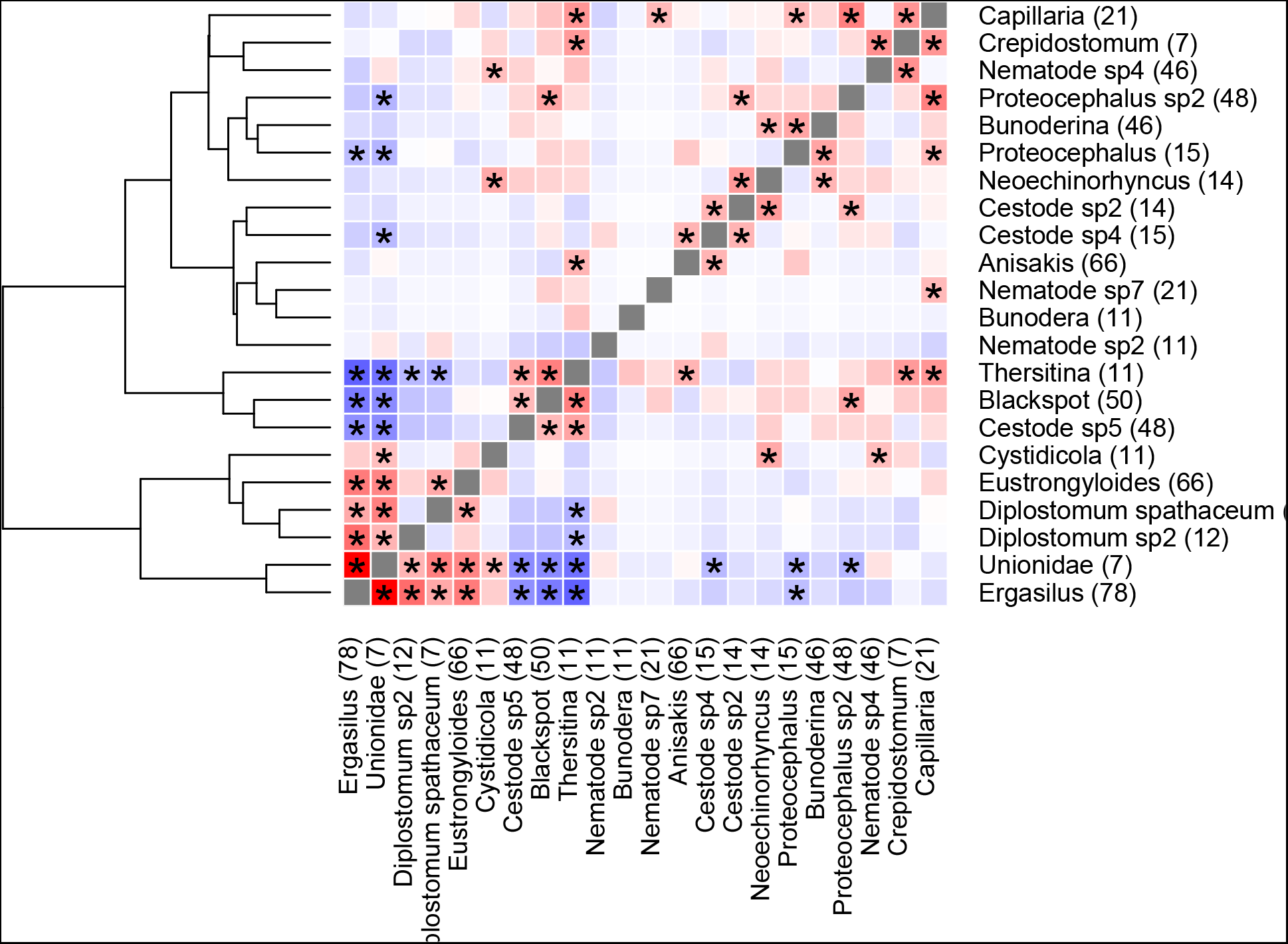
Co-occurrence matrix of parasites among individual stickleback in Roberts lake (N = 250 fish). Numbers represent the number of hosts carrying a given parasite.

**Figure S3.**
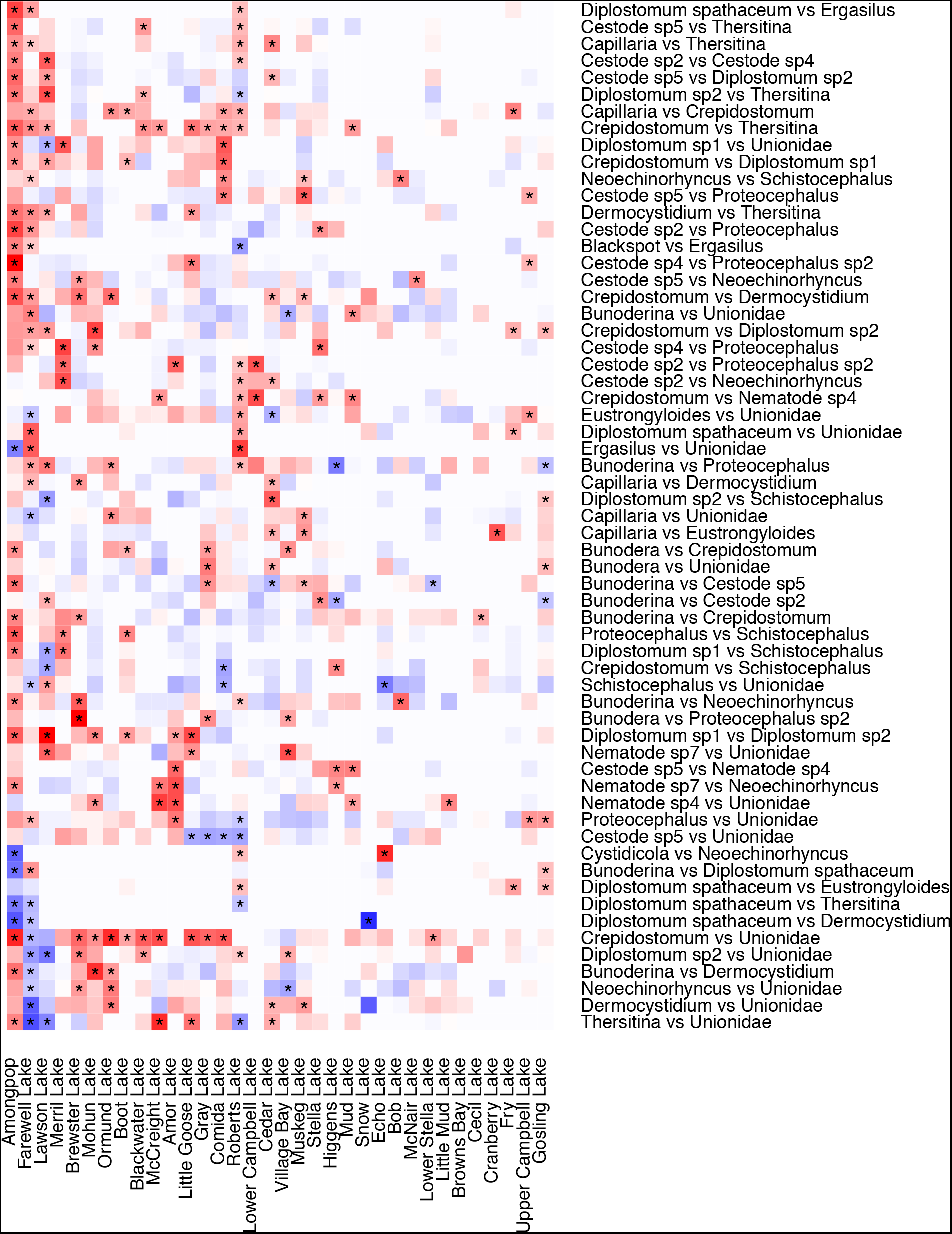
Parasite-parasite correlations are inconsistent among host populations. Each column is a host population, with rows representing parasite-parasite comparisons. The farthest-left column represents the among-population correlation matrix (Fig. 5), for comparison. Red and blue denote positive and negative correlations, respectively. White indicates pairwise comparisons for which one or both parasites are absent in a given lake. Asterisks indicate significant comparisons within each population after within-population FDR correction.

**Figure S4.**
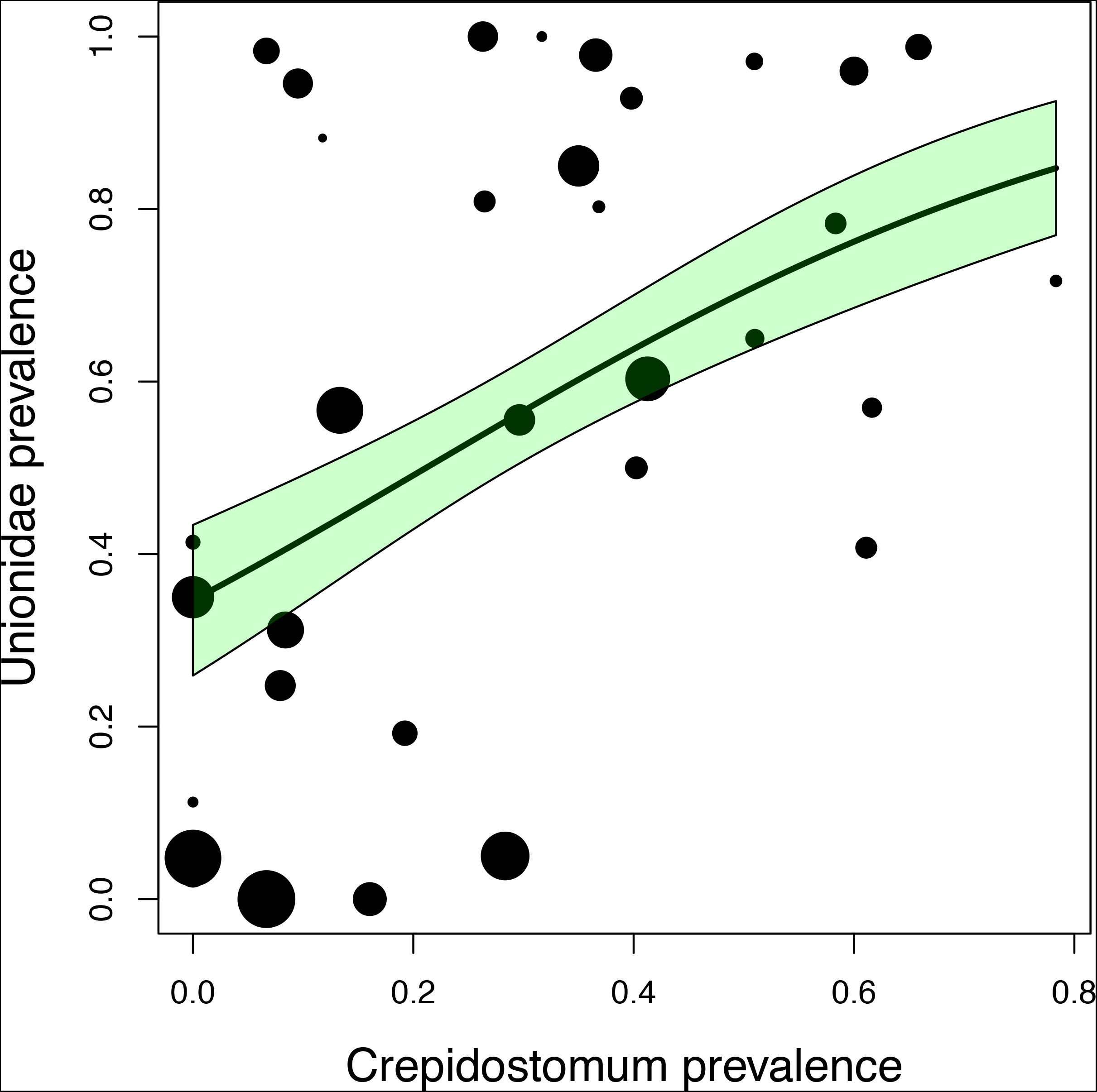
An example of parasite-parasite co-occurrence among lakes. Each point is a population (points scaled by their sample size), and the trendline is a Poisson general linear model curve fit with a one standard error confidence interval.

**Figure S5.**
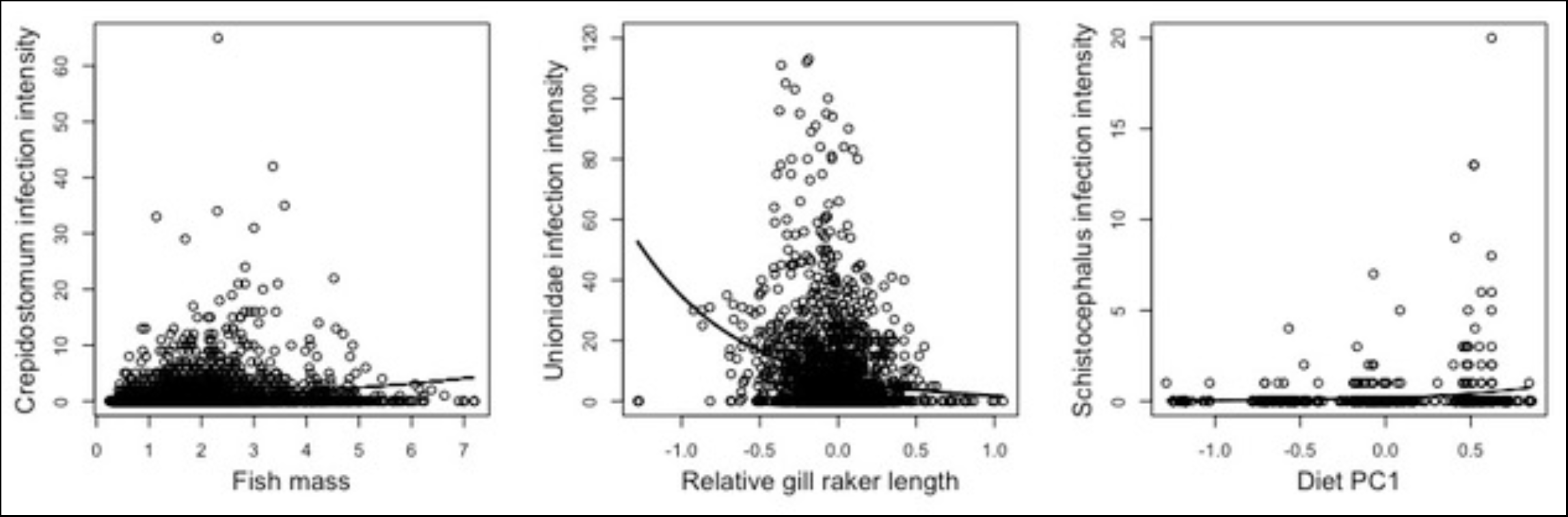
Examples of associations between parasite infection intensity and host individual’s traits, for (A) mass, (B) trophic morphology, and (C) diet, pooled across all individuals. Statistical details of these associations, and other trait and parasite combinations, are summarized in Table S1.

**Figure S6.**
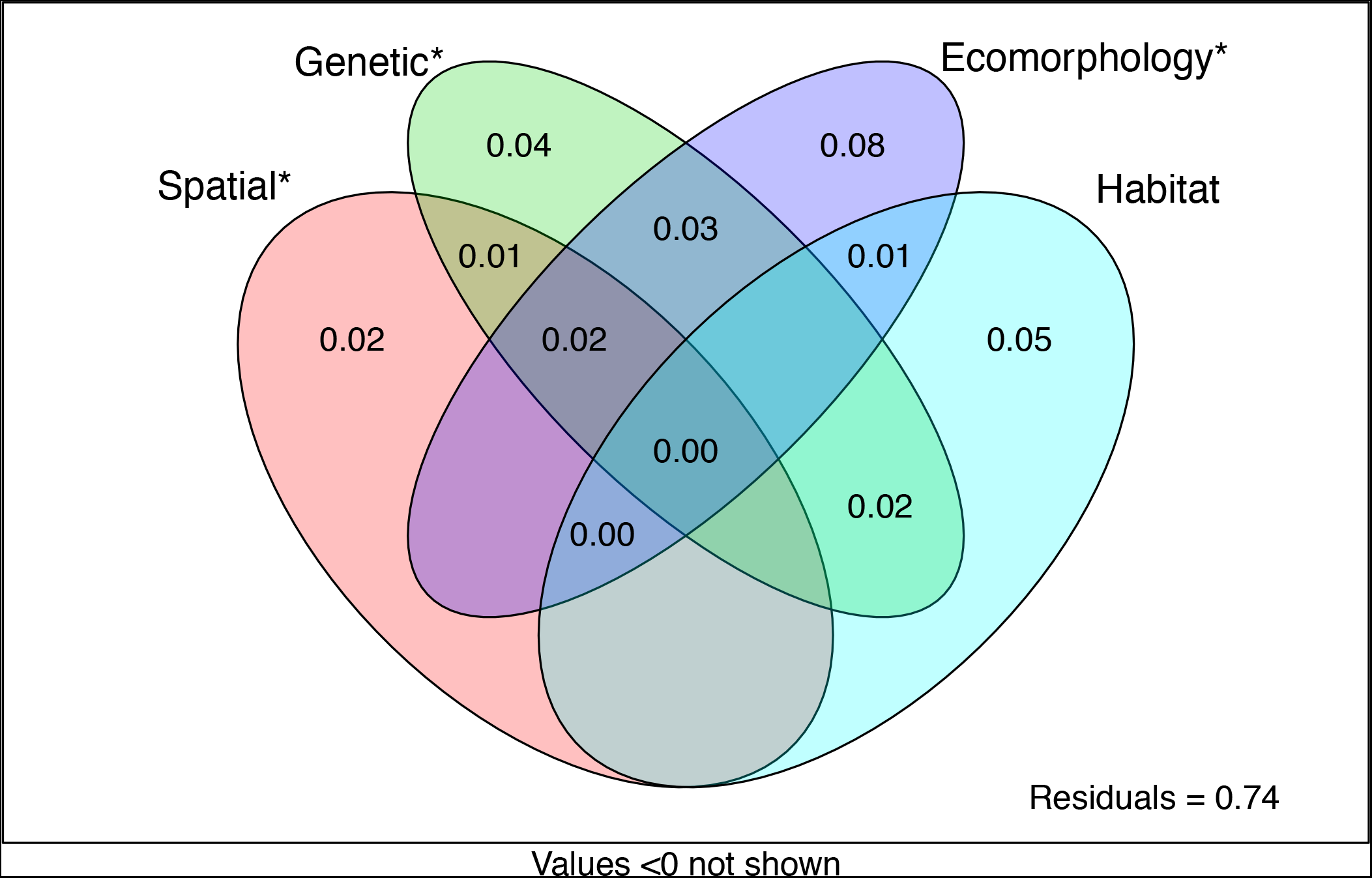
Variance partitioning analysis, estimating the extent to which among-lake variation in parasite community composition depends on spatial distance, genetic distance, phenotypic distance, and habitat differences. Numbers in each cell represent represent the percent variance explained by each factor or interactions among factors (at intersections).

**Figure S7.**
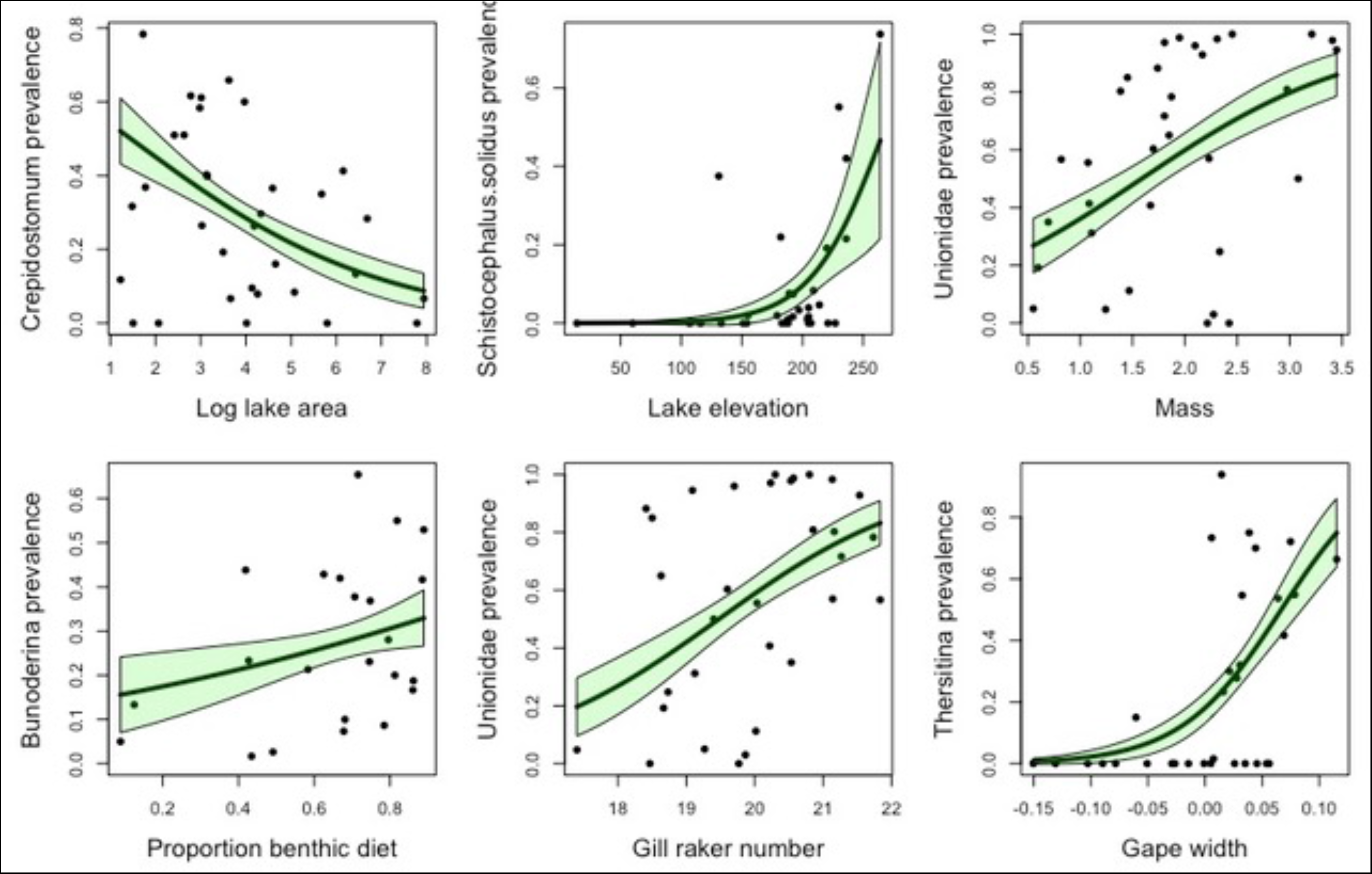
Examples of associations between parasite prevalence (% hosts infected) and lake or host population traits, for (A) lake area, (B) lake elevation, (C) mean fish mass, (D) mean proportion benthic prey, (E) mean gill raker number, and (F) mean gape width. GLM trendlines with one standard error confidence estimates are plotted. Lakes are the level of replication in this figure. These associations are summarized in Table S2.

**Figure S8.**
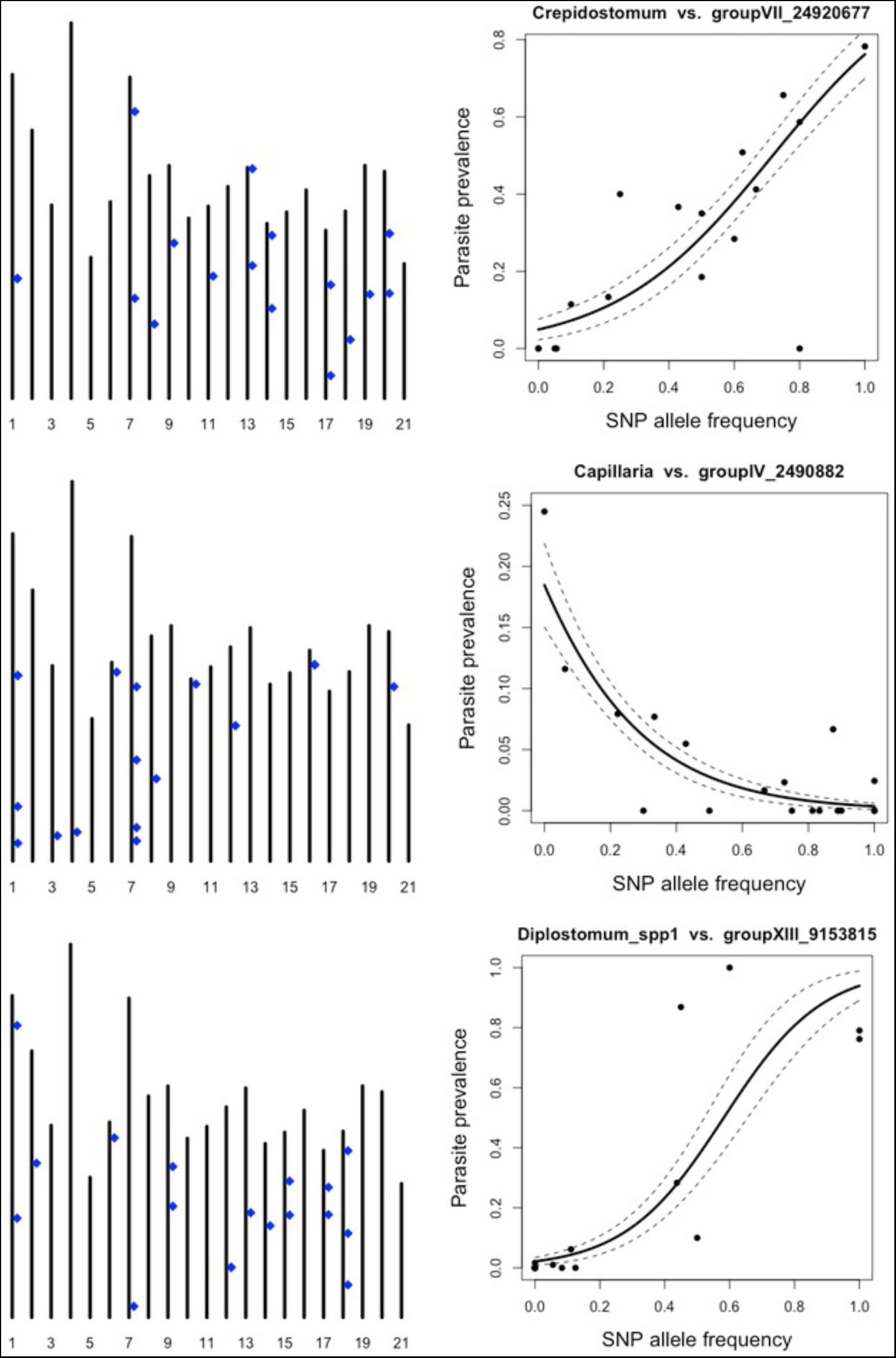
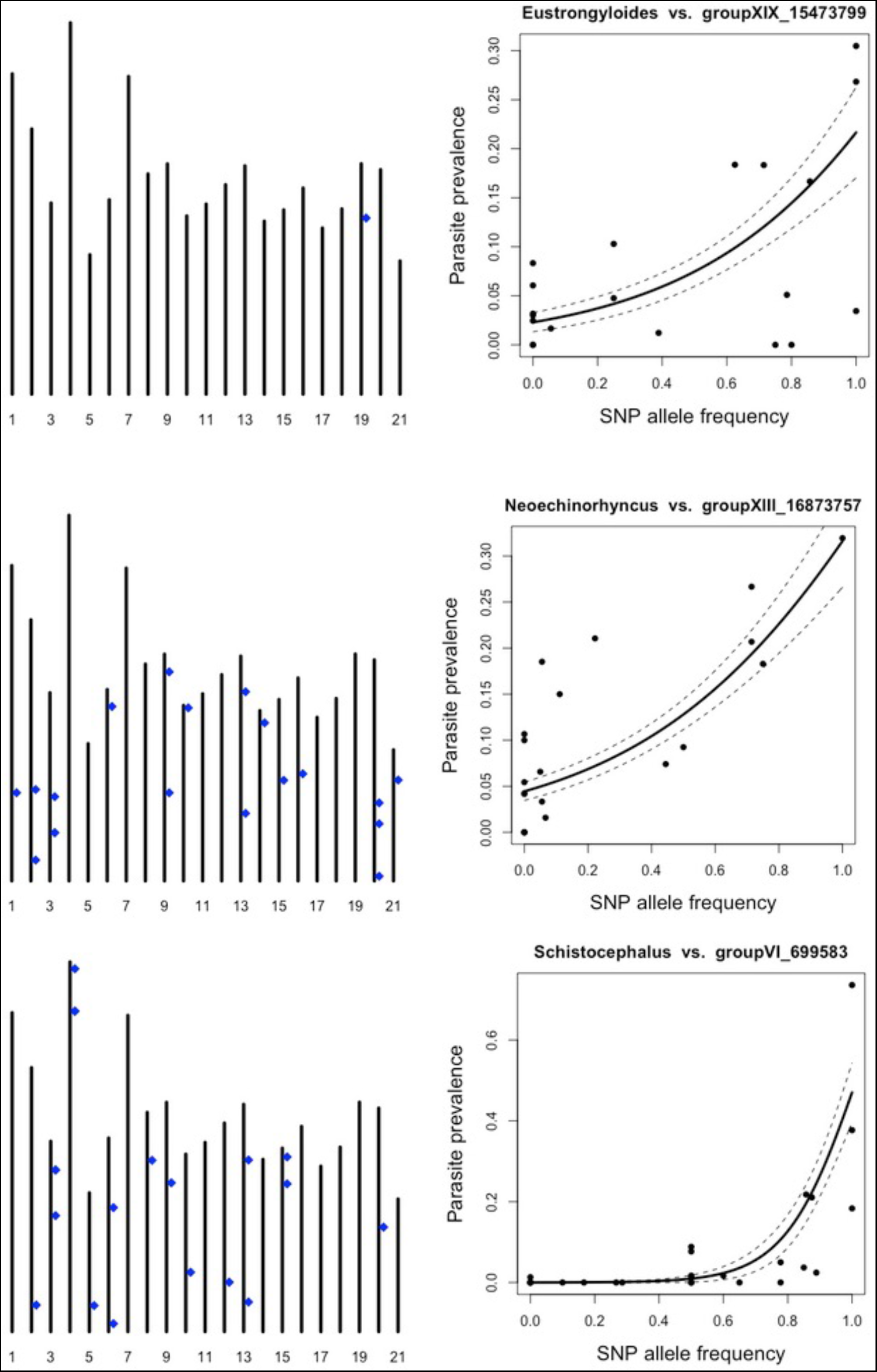
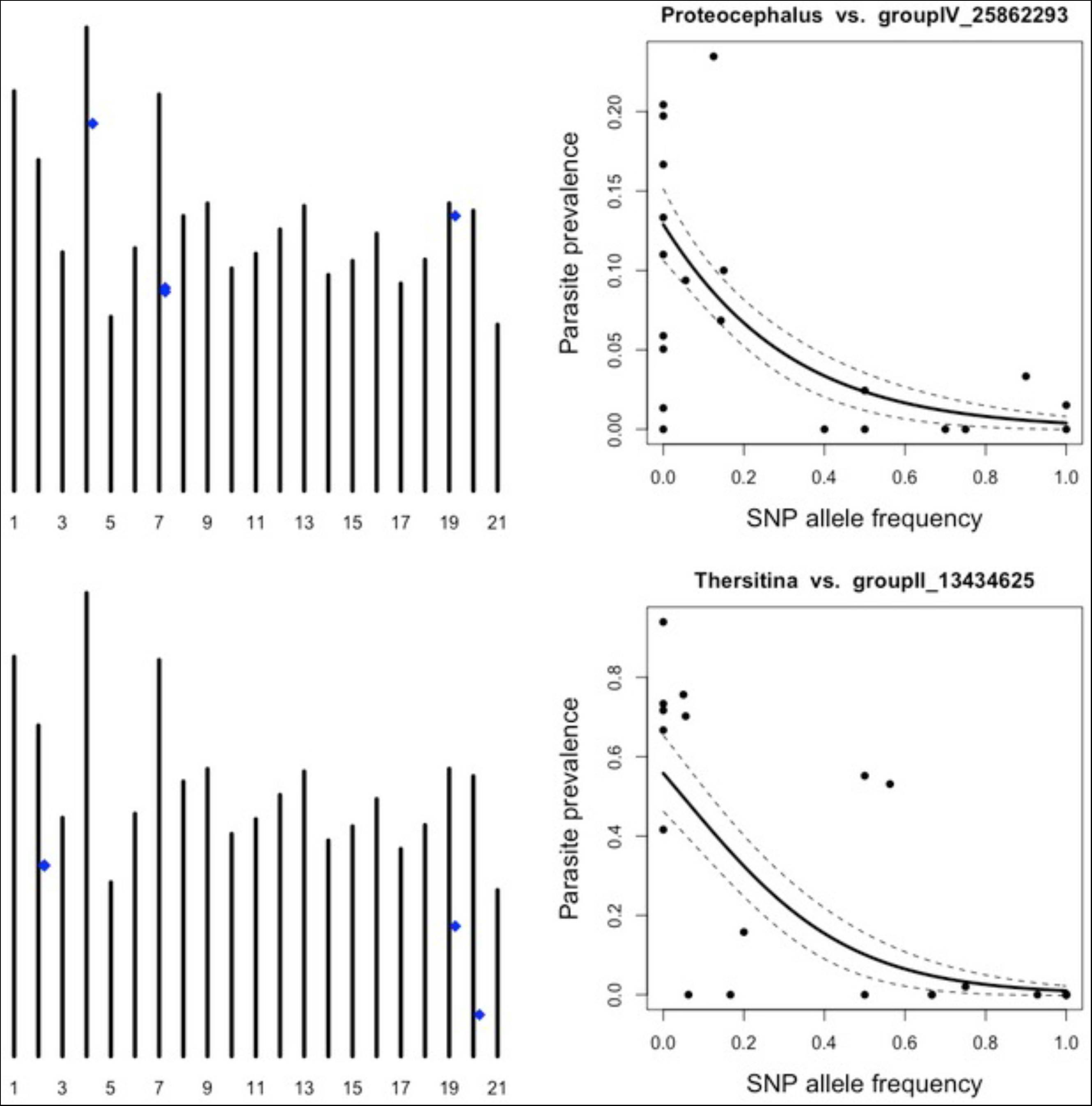
Example of strong GWAS results for various parasite taxa, chosen as illustrative examples. For each taxon we plot the position of GWAS associations (blue dots) along chromosomes, showing only the top 20 associations significant after Bonferroni correction. Next to each map, we plot the association between SNP frequency versus the parasite prevalence, for one of the strongest-effect SNPs for that parasite. Points represent populations, with a binomial GLM curve fit. Above each association plot we list the parasite taxon and the linkage group and base position on that chromosome.

**Figure S9.**
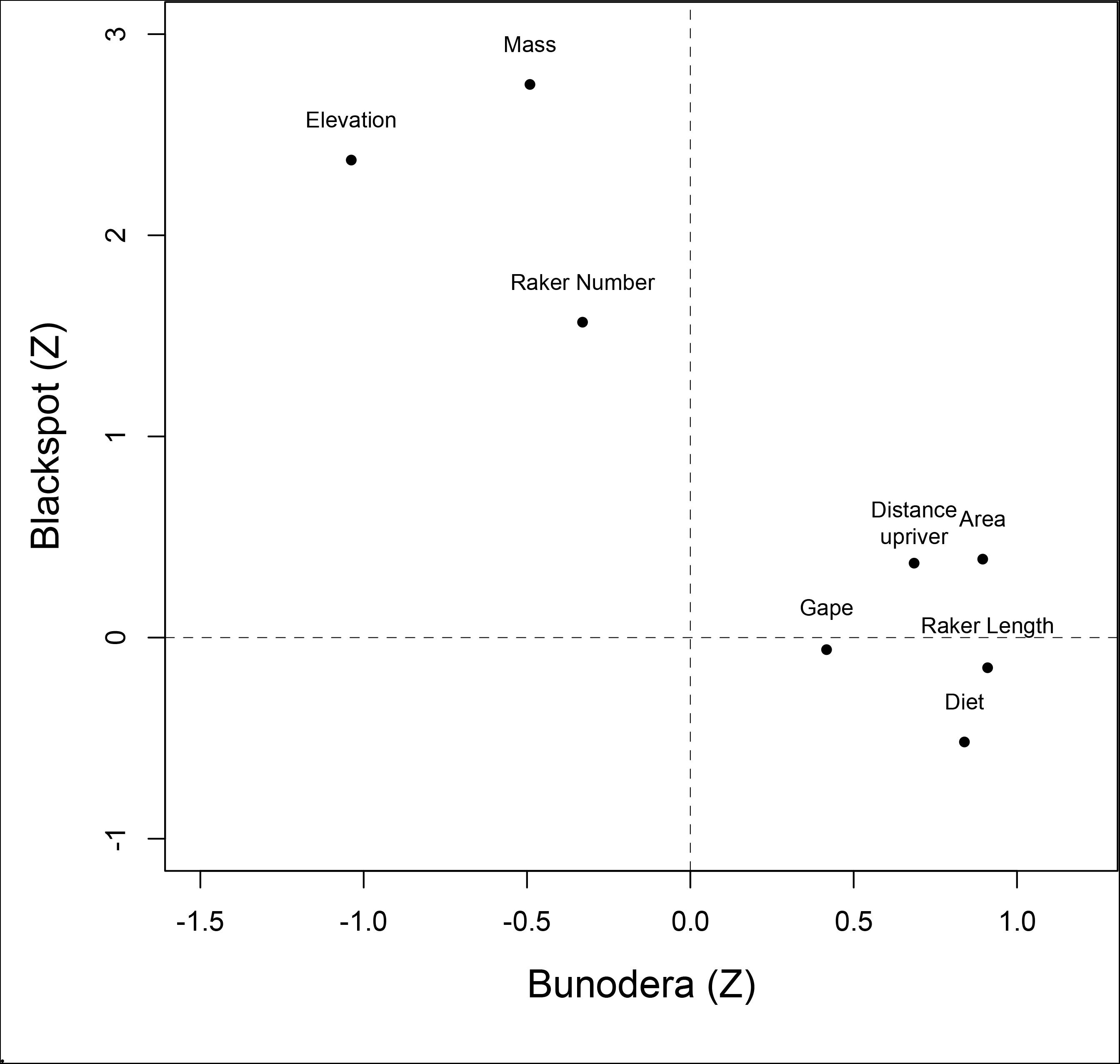
An example of negative co-dependence (Fig. 5C). Bunodera and Blackspot exhibit opposing associations with host population trait means, and lake characteristics, at the scale of among-lake variation.

**Table S1.** Summary of sampled populations, locations, sample sizes, and mean parasite taxon richness.

| Site name | Longitude | Latitude | Total | Morphology | Diet | ddRAD | Habitat | Watershed |
| --- | --- | --- | --- | --- | --- | --- | --- | --- |
| Adam and Eve Estuary | -131.9426 | 50.4012 | 99 | 99 | 0 | 10 | Estuary | Adam River |
| Amor Bridge | -125.6639 | 50.2543 | 60 | 30 | 23 | 10 | Stream | Amor River |
| Amor Lake | -125.5791 | 50.1581 | 60 | 30 | 30 | 10 | Lake | Amor River |
| Blackwater Lake | -125.5878 | 50.1794 | 82 | 30 | 25 | 16 | Lake | Amor River |
| Bob Lake | -125.5281 | 50.3045 | 80 | 61 | 61 | 10 | Lake | Pye River |
| Boot Lake | -125.5270 | 50.0467 | 93 | 93 | 0 | 0 | Lake | Campbell River |
| Brewster Lake | -125.5779 | 50.0924 | 63 | 30 | 28 | 10 | Lake | Campbell River |
| Browns Bay Lake | -125.4159 | 50.1580 | 29 | 0 | 0 | 10 | Lake | Browns Bay Creek |
| Campbell River Marsh | -125.2570 | 50.0387 | 105 | 105 | 0 | 0 | Estuary | Campbell River |
| Campbell River Point | -125.2574 | 50.0476 | 173 | 173 | 0 | 0 | Estuary | Campbell River |
| Cecil Lake | -125.5440 | 50.2361 | 100 | 100 | 0 | 10 | Lake | Amor River |
| Cedar Lake | -125.5664 | 50.2029 | 98 | 30 | 29 | 10 | Lake | Amor River |
| Comida Lake | -125.5283 | 50.1502 | 86 | 86 | 0 | 0 | Lake | Mohun River |
| Comida Stream | -125.5215 | 50.1339 | 192 | 192 | 0 | 0 | Stream | Mohun River |
| Cranberry Lake | -125.4553 | 50.0883 | 66 | 66 | 0 | 10 | Lake | Mohun River |
| Echo Lake | -125.4084 | 49.9848 | 68 | 68 | 0 | 10 | Lake | Quinsam River |
| Farewell Lake | -125.5905 | 50.2013 | 216 | 216 | 0 | 0 | Lake | Amor River |
| Farewell Stream | -125.5992 | 50.2058 | 260 | 260 | 0 | 0 | Stream | Amor River |
| Fry Lake | -125.5720 | 50.0244 | 101 | 101 | 0 | 11 | Lake | Campbell River |
| Gosling Lake | -125.5023 | 50.0483 | 147 | 146 | 0 | 0 | Lake | Campbell River |
| Gray Lake | -125.5953 | 50.0553 | 75 | 30 | 27 | 9 | Lake | Campbell River |
| Higgins Lake | -125.5088 | 50.0680 | 81 | 30 | 29 | 10 | Lake | Campbell River |
| Lawson Lake | -125.5790 | 50.0362 | 82 | 30 | 27 | 10 | Lake | Campbell River |
| Little Goose Lake | -125.4890 | 50.1634 | 76 | 31 | 28 | 10 | Lake | Mohun River |
| Little Mud Lake | -125.5505 | 50.2069 | 60 | 30 | 29 | 10 | Lake | Amor River |
| Lower Campbell Lake | -125.4760 | 50.0338 | 30 | 30 | 30 | 10 | Lake | Campbell River |
| Lower Stella Lake | -125.5501 | 50.3107 | 104 | 30 | 30 | 10 | Lake | Pye River |
| McCreight Lake | -125.6390 | 50.3295 | 60 | 30 | 28 | 10 | Lake | Amor River |
| McCreight Stream | -125.6457 | 50.3321 | 60 | 30 | 29 | 10 | Stream | Amor River |
| McNair Lake | -125.5744 | 50.2281 | 60 | 30 | 28 | 10 | Lake | Amor River |
| Merril Lake | -125.5568 | 50.0593 | 38 | 30 | 28 | 10 | Lake | Campbell River |
| Mohun Creek | -125.3767 | 50.1113 | 27 | 27 | 21 | 10 | Stream | Mohun River |
| Mohun Lake | -125.4890 | 50.1640 | 60 | 30 | 28 | 10 | Lake | Mohun River |
| Mud Lake | -125.5541 | 50.1871 | 60 | 30 | 28 | 10 | Lake | Amor River |
| Muskeg Lake | -125.5808 | 50.1989 | 104 | 30 | 28 | 10 | Lake | Amor River |
| Ormund Lake | -125.5267 | 50.1795 | 60 | 30 | 28 | 10 | Lake | Amor River |
| Pye Creek | -125.5772 | 50.3122 | 60 | 29 | 27 | 0 | Stream | Pye River |
| Pye Outlet | -125.5191 | 50.3345 | 73 | 73 | 0 | 10 | Estuary | Pye River |
| Roberts Lake | -125.5428 | 50.2158 | 250 | 250 | 0 | 0 | Lake | Amor River |
| Roberts Stream | -125.5681 | 50.2339 | 354 | 345 | 0 | 0 | Stream | Amor River |
| Sayward Estuary | -125.9464 | 50.3769 | 100 | 100 | 0 | 10 | Estuary | Salmon River |
| Snow Lake | -125.5937 | 50.3127 | 17 | 17 | 3 | 9 | Lake | Pye River |
| Stella Lake | -125.5175 | 50.2992 | 60 | 30 | 30 | 11 | Lake | Pye River |
| Upper Campbell Lake | -125.5841 | 49.9953 | 105 | 30 | 29 | 10 | Lake | Campbell River |
| Village Bay Lake | -125.1878 | 50.1708 | 81 | 60 | 59 | 0 | Lake | Village Bay |
| Village Bay Stream | -125.2064 | 50.1675 | 60 | 30 | 29 | 0 | Stream | Village Bay |

**Table S2.**
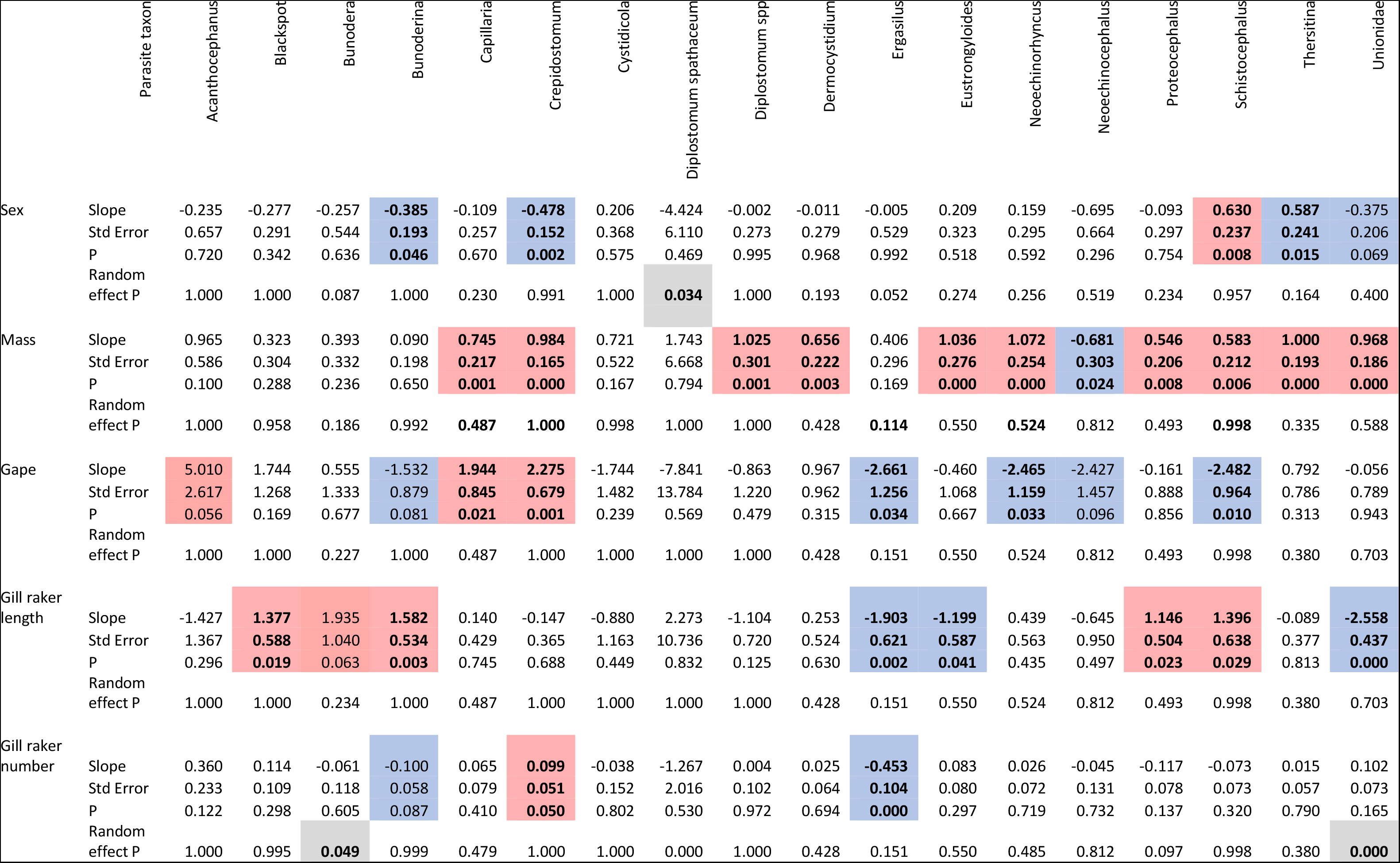
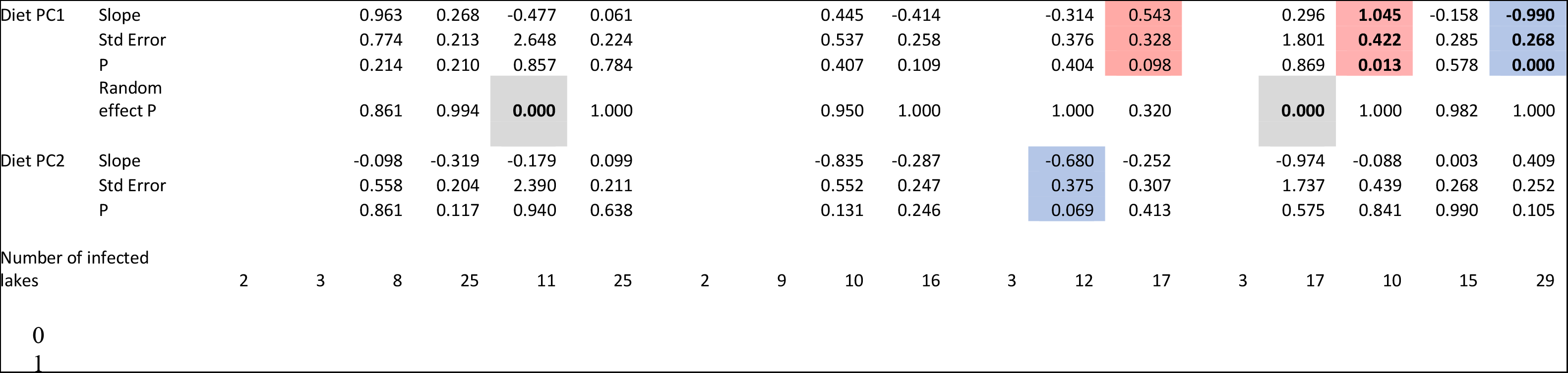
(begins on next page). General linear model results from individual-host scale regressions of parasite intensity on individual host traits. We provide slope estimates and standard errors and P-values for the main effects listed, and a P-value for the random slope effect of population. Positive and negative effects are shaded red and blue, respectively, for all significant (P < 0.05, also in bold font) and marginally significant (P < 0.1, not bold font) associations. Significant random slope effects of lake are shaded grey.

**Table S3.**
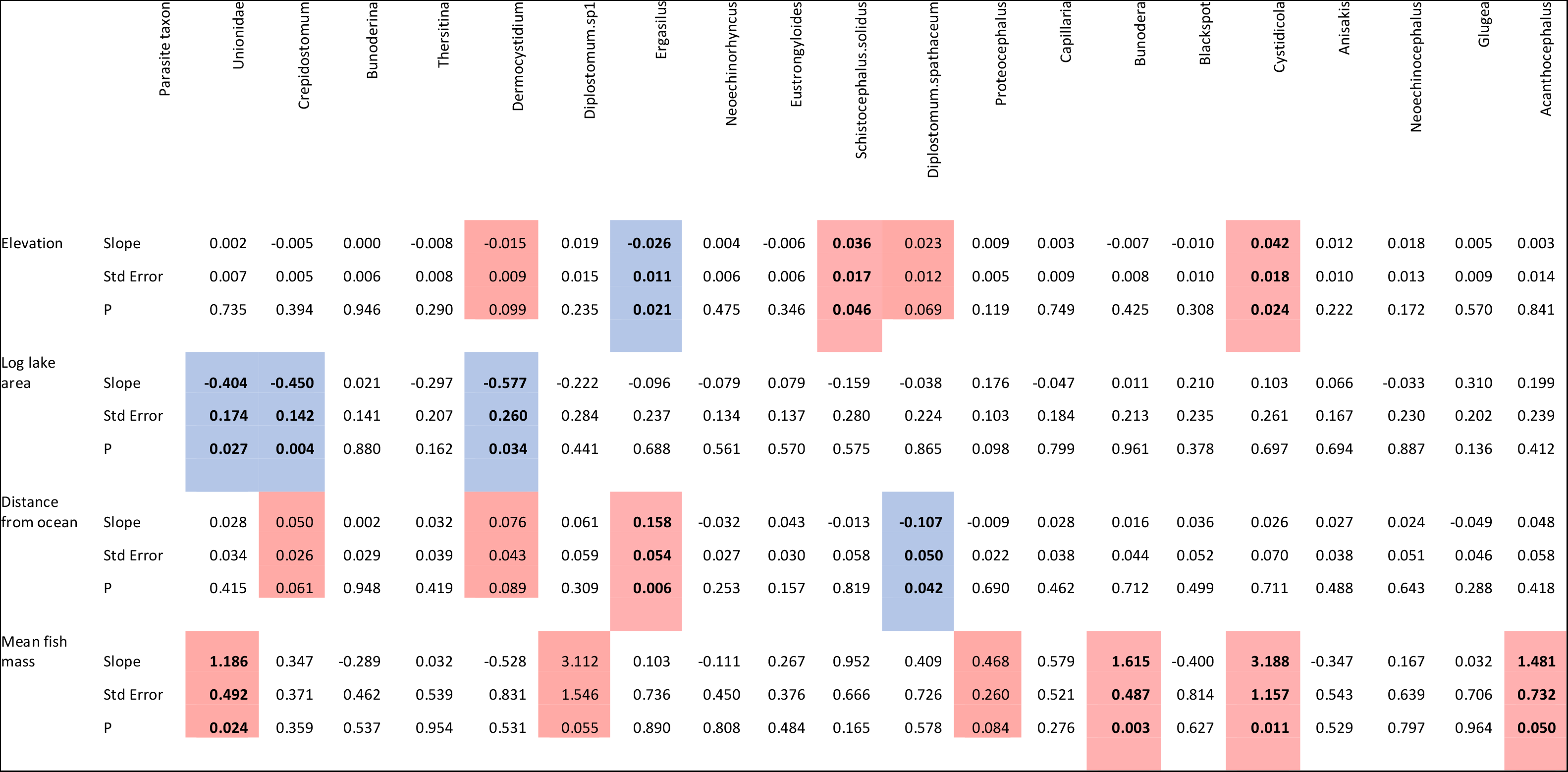
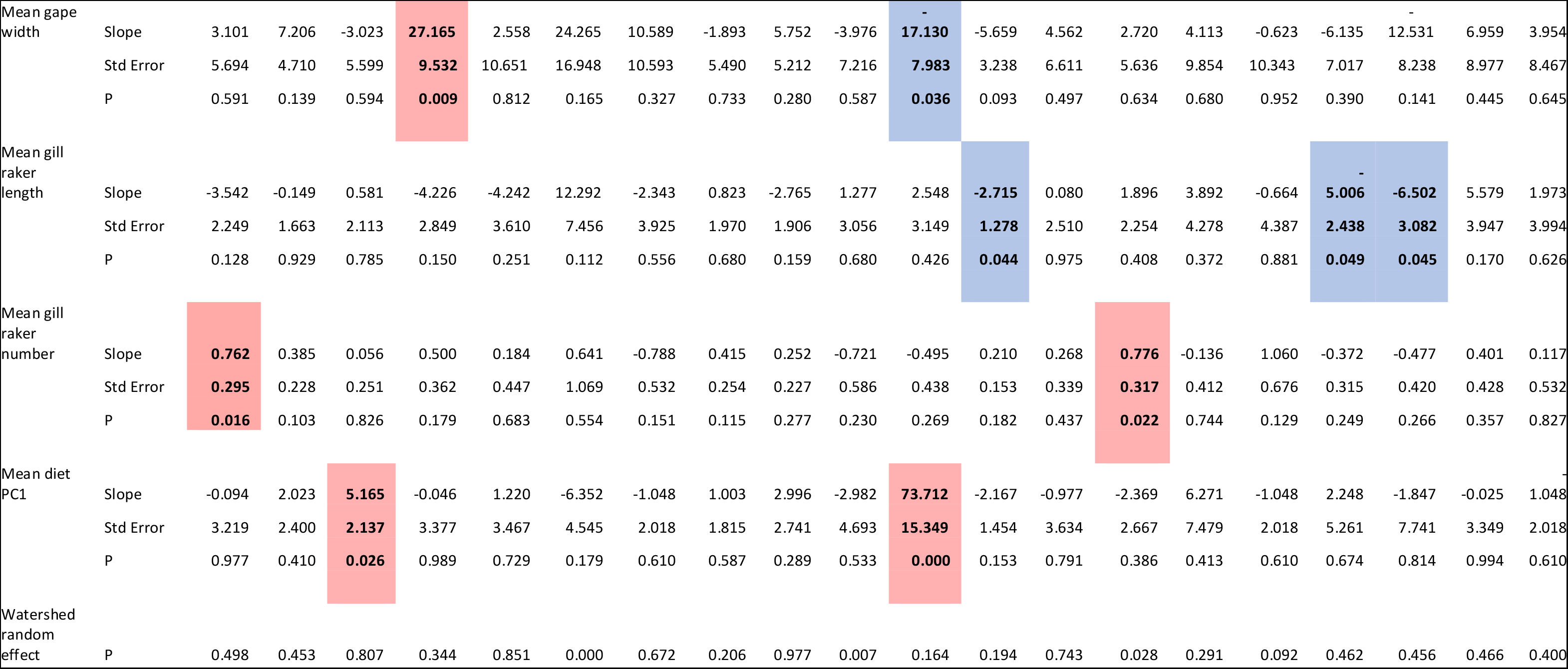
General linear model results from host-population scale regressions of parasite prevalence on lake and population traits (e.g., trait means). We provide slope estimates and standard errors and P-values for the main effects listed. Positive and negative effects are shaded red and blue, respectively, for all significant (P < 0.05, also in bold font) and marginally significant (P < 0.1, not bold font) associations. Parasite taxa (columns) are ordered by average abundance across the entire metacommunity from most abundant (left) to least (right), 7

